# Interrogation and validation of the interactome of neuronal Munc18-interacting Mint proteins with AlphaFold2

**DOI:** 10.1101/2023.02.20.529329

**Authors:** Saroja Weeratunga, Rachel S. Gormal, Meihan Liu, Denaye Eldershaw, Emma K. Livingstone, Anusha Malapaka, Tristan P. Wallis, Adekunle T. Bademosi, Anmin Jiang, Michael D. Healy, Frederic A. Meunier, Brett M. Collins

**Author notes:** Corresponding author: Brett Collins.

## Abstract

Munc18-interacting proteins (Mints) are multi-domain adaptors that regulate neuronal membrane trafficking, signalling and neurotransmission. Mint1 and Mint2 are highly expressed in the brain with overlapping roles in the regulation of synaptic vesicle fusion required for neurotransmitter release by interacting with the essential synaptic protein Munc18-1. Here, we have used AlphaFold2 to identify and then validate the mechanisms that underpin both the specific interactions of neuronal Mint proteins with Munc18-1 as well as their wider interactome. We find a short acidic α-helical motif (AHM) within Mint1 and Mint2 is necessary and sufficient for specific binding to Munc18-1 and binds a conserved surface on Munc18-1 domain3b. In Munc18-1/2 double knockout neurosecretory cells mutation of the Mint-binding site reduces the ability of Munc18-1 to rescue exocytosis, and although Munc18-1 can interact with Mint and Sx1a proteins simultaneously *in vitro* we find they have mutually reduced affinities, suggesting an allosteric coupling between the proteins. Using AlphaFold2 to then examine the entire cellular network of putative Mint interactors provides a structural model for their assembly with a variety of known and novel regulatory and cargo proteins including ARF3/ARF4 small GTPases, and the AP3 clathrin adaptor complex. Validation of Mint1 interaction with a new predicted binder TJAP1 provides experimental support that AlphaFold2 can correctly predict interactions across such large-scale datasets. Overall, our data provides insights into the diversity of interactions mediated by the Mint family and shows that Mints may help facilitate a key trigger point in SNARE complex assembly and vesicle fusion.

## INTRODUCTION

Synaptic vesicle fusion and release of neurotransmitters requires the formation of specific complexes between vesicular and plasma membrane SNARE proteins (soluble *<u>N</u>*-ethylmaleimide-sensitive factor <u>a</u>ttachment <u>re</u>ceptors). The vesicular v-SNAREs and target t-SNAREs form an α-helical coiled-coil assembly that provides the necessary energy to bring the two opposing lipid membranes together for membrane fusion. Munc18-1 (also called Munc18a, syntaxin-binding protein 1 (STXBP1), and neuronal Sec1 (nSec1)) is a member of the Sec1/Munc18 (SM) protein family, and an essential regulatory protein required for the assembly of this SNARE complex in neuronal membrane fusion. Munc18-1 binds with high affinity to the target or Qa-SNARE Syntaxin1a (Sx1a) and mediates both its trafficking and its incorporation into the SNARE complex with the Qbc-SNARE SNAP25 and the vesicle or R-SNARE Vamp2.

Importantly, Munc18-1 also interacts with other regulatory proteins including Munc13 and the two Munc-interacting protein (Mint) paralogues Mint1 and Mint2. Mints (also known as X11, amyloid precursor protein-binding family A (APBA) or Lin-10 proteins) are multi domain scaffolds that participate in a host of protein-protein interactions. The human genome encodes three Mint homologues, Mint1, Mint2 and Mint3 (Rogelj *et al*, 2006) (**Fig. 1A**). Mint1 and Mint2 are highly enriched in brain and spinal cord tissue, while Mint3 is ubiquitously expressed (Biederer *et al*, 2002; Ho *et al*, 2002b; Lee *et al*, 2003; Motodate *et al*, 2016; Tomita *et al*, 1999). Structurally, Mint proteins are composed of conserved C-terminal phosphotyrosine binding (PTB) and tandem PSD95/Dlg/ZO- 1 (PDZ) domains and possess highly extended intrinsically disordered N-terminal domains that mediate paralogue-specific interactions. As scaffold proteins, the PTB domains of the Mint proteins interact with various ligands containing NPxY sequences (x is any amino acid) such as the amyloid precursor protein (APP), APP-like proteins APLP1 and APLP2, and TrkA (Borg *et al*, 1996; Borg *et al*, 1998; Caster & Kahn, 2013; Chaufty *et al*, 2012; Ho *et al*, 2008; Matos *et al*, 2012; Saito *et al*, 2011; Sakuma *et al*, 2009; Sullivan *et al*, 2014; Tomita *et al*., 1999; Xie *et al*, 2013; Zhang *et al*, 2009; Zhang *et al*, 1997), while the PDZ domains of the Mint proteins bind to C-terminal PDZ binding motifs (PDZbms) in molecules including NMDA receptors, kalirin-7, neurexins, ApoER2 and LDLR (Biederer & Sudhof, 2000; Gotthardt *et al*, 2000; Jones *et al*, 2014; Minami *et al*, 2010; Motodate *et al*, 2019). The N-termini of the Mint proteins contain sequences that mediate both conserved and isoform specific protein-protein interactions; for example, an N-terminal sequence specific to Mint1 called the CASK-interacting domain (CID) binds specifically to Ca^2+^/calmodulin-dependent serine protein kinase (CASK/Lin-2) (a multi domain adapter and Mg^2+^-independent S/T kinase) forming a tripartite complex with Veli/Lin-7 family proteins and neurexins at the pre-synapse (Borg *et al*., 1998; Stafford *et al*, 2011; Tabuchi *et al*, 2002; Wu *et al*, 2020; Zhang *et al*, 2020). In contrast, the N-terminal regions of Mint1 and Mint2 proteins can both interact with Munc18-1 via a sequence termed the Munc18-1-interacting domain (MID) (Biederer & Sudhof, 2000; Graham *et al*, 2011; Han *et al*, 2014; Ho *et al*., 2002b; Okamoto & Sudhof, 1997, 1998; Park *et al*, 2012). However, until recently the mechanism of Munc18-1 interaction was unknown.

**Figure 1.**
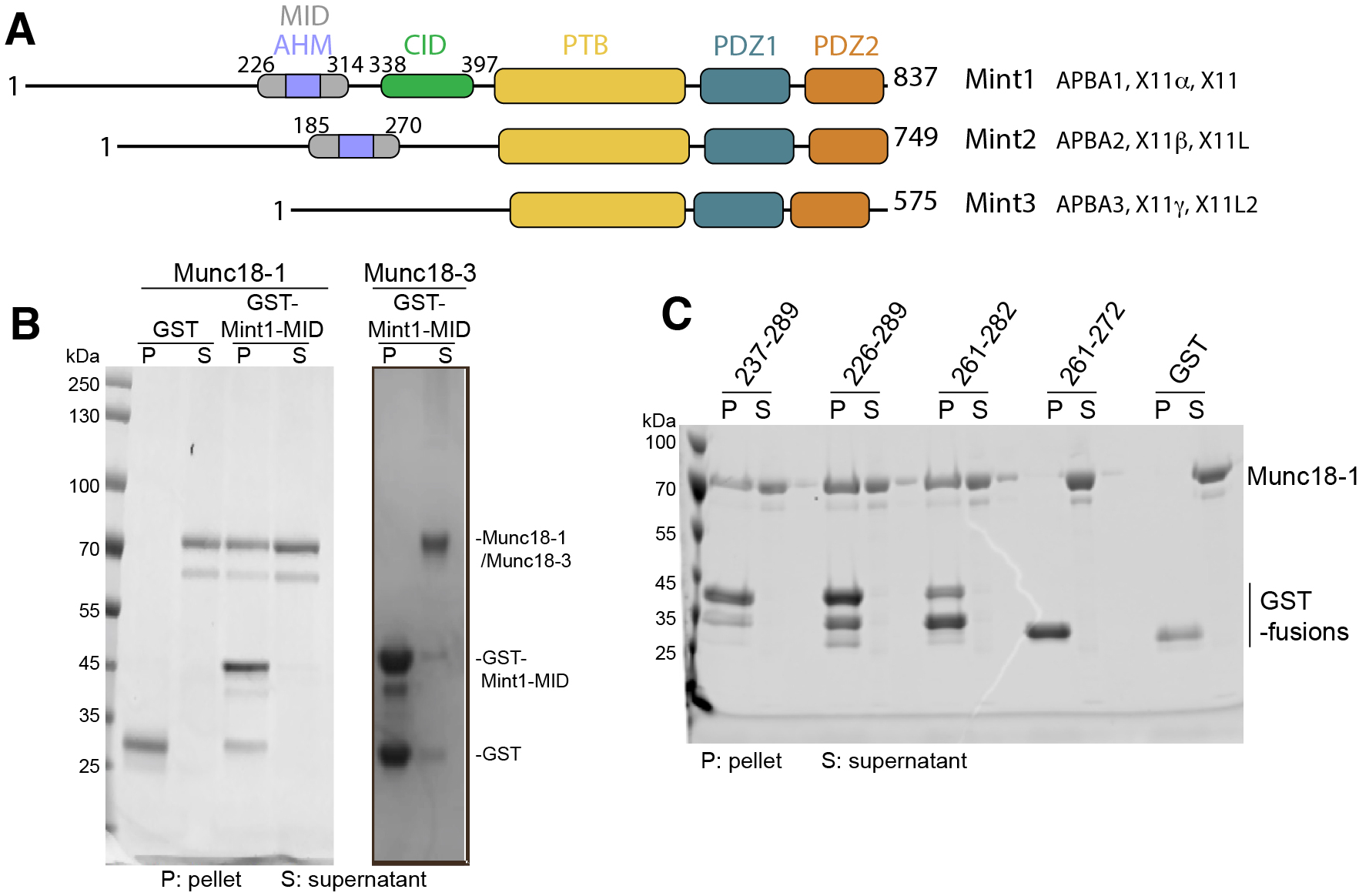
The Mint1 MID interacts directly with Munc18-1 but not Munc18-3. **(A)** Schematic diagram of the human Mint proteins. (AHM, acidic α-heilcal motif; MID, Munc18-1- interacting domain; CID, CASK interacting domain; PTB, phosphotyrosine binding domain; PDZ1 and PDZ2, Psd95/Dlg/ZO1 domains). **(B)** Pulldowns with GST-Mint1 MID show a direct interaction with purified Munc18-1 but not Munc18-3. Image shows Coomassie Blue stained gel. **(C)** Pulldowns with GST-tagged Mint1 truncated sequences identify residues 261-282 as sufficient and required for Munc18-1 binding. Image shows Coomassie Blue stained reducing SDS-PAGE gels.

The importance of Mint proteins in neuronal function is apparent in studies showing homozygous deletions of Mint1 or Mint2 display disrupted GABAergic transmission (Ho *et al*, 2003; Sano *et al*, 2006), while knockout of both Mint1 and Mint2 neuronal isoforms leads to lethality at birth in most animals, with surviving mice displaying deficits in motor behaviours and spontaneous neurotransmitter release. A prominent phenotype of knockout animals is an alteration in APP trafficking and processing to the Alzheimer’s plaque peptide amyloid β (Aβ) (Gross *et al*, 2013; Sullivan *et al*., 2014), although the changes appear to be dependent on specific knockout conditions with both an increase (Kondo *et al*, 2010; Saluja *et al*, 2009; Sano *et al*., 2006) and decrease (Ho *et al*., 2008; Sullivan *et al*., 2014) in Aβ production reported. Thus, the molecular interactions between Mints, synaptic proteins and trafficking partners involved in neurodegenerative diseases is of broad interest to the field.

Mutations in the core neuronal pre-synaptic machinery including Munc18-1 and Sx1a lead to a distinct but overlapping set of neurodevelopmental disorders, with symptoms including neurodevelopmental delay, intellectual disability and early infantile epileptic encephalopathy (Lanoue *et al*, 2019; Verhage & Sorensen, 2020). Mutations in the Mint1-assocated CASK protein have also been found to cause neurodevelopmental disorders with similar features, including mental retardation and microcephaly with ophthalmic atrophy (Hayashi *et al*, 2017; Hsueh, 2009; LaConte *et al*, 2019; Piluso *et al*, 2009), and the human disease linked missense variant p.Leu209Pro in CASK linked to optic nerve hypoplasia specifically disrupts Mint1 binding without impacting other protein interactions (LaConte *et al*., 2019). Mutations in the Mint proteins themselves have not yet been specifically linked to similar disorders, although copy number variations in Mint2 have been described in patients with epilepsy and intellectual disability (Peycheva *et al*, 2018). In addition, variants in Mint2 have been linked to autism spectrum disorder, potentially through disruption of neurexin trafficking (Babatz *et al*, 2009; Lin *et al*, 2019). By assessing the mechanisms responsible for Mint1 binding to a range of protein partners, we aim to provide a framework that can contribute to the understanding of how Mint1 might modulate a role during neurotransmission and trafficking.

Here we explored the use of AlphaFold2 to identify and predict the structural basis for Mint protein interactions across their putative interactome. We initially focused on the interaction with SNARE regulatory protein Munc18-1, where AlphaFold2 confidently predicts a direct association with Mint1 and Mint2, and extensively validate the mechanism of binding between the Mint MID domain and a a novel binding site in domain 3b of Munc18-1. We define a minimal short linear interaction motif (SLiM) of thirteen amino acids in Mint1 and Mint2 that is required and sufficient for Munc18-1 interaction. The AlphaFold2 derived model of the Mint1 and Mint2 sequences bound to Munc18-1 shows that the peptides form an acidic α-helical motif (AHM) bound to Munc18-1 domain 3b. Providing further evidence for the ability of AlphaFold2 to accurately predict novel protein-peptide complexes, while this was predicted and validated prior to any experimental structure, recent work by Li and colleagues described a crystal structure revealing an essentially identical association (Li *et al*, 2023). Munc18-1 mutation R388A blocks Mint-1 interaction *in vitro*, and significantly reduced the number of exocytic events detected using VAMP2-pHluorin unquenching in Munc18-1/2 double knockout PC12 cells. Although the respective Mint1/2 and Sx1a binding sites in domain 3b and domains 1/3a are distinct from each other, their individual Munc18-1 *in vitro* binding affinities are reduced in the presence of the other ligand. We speculate that this antagonistic allosteric interaction between the two proteins may be important for regulating Munc18-1 templating of SNARE-complex formation. Building on the successful structural predictions of Mint1 and Munc18-1 with AlphaFold2, we have explored the wider network of interactions mediated by the Mint scaffolds. We confidently identify a novel binding site in the Mint PDZ domains for ARF small GTPases, and experimentally validate a non-canonical site in the Mint PTB domain, distinct from the NPxY peptide motif-binding site, that mediates binding to tight junction-associated protein 1 (TJAP1). These studies provide an overall model for neuronal Mint1 and Mint2 assembly with both effector and regulatory proteins.

## RESULTS

### Mapping the Munc18-1-binding sequence of Mint1 and Mint2

A number of studies have shown that both Mint1 and Mint2 homologues can interact with Munc18- 1, using methods including immunoprecipitation and yeast two-hybrid assays (Biederer & Sudhof, 2000; Ho *et al*, 2002a; Lee *et al*, 2004; M. Okamoto, 1997; Okamoto & Sudhof, 1997). Mint proteins are multi-domain scaffolds with a C-terminal PTB domain and tandem PDZ domains preceded by a long and unfolded N-terminal sequence that shows low sequence homology across the family (**Fig. 1A; Fig. S1**). Munc18-1 binding has been mapped by yeast two-hybrid assays to a region termed the ‘Munc18-interacting domain’ (MID) within the N-terminus of the neuronal Mint1 and Mint2 proteins consisting of residues 226-314 and 185-270 respectively (Okamoto & Sudhof, 1997). We first confirmed the direct interaction of Mint1 with purified Munc18-1, performing a GST pull-down assay using the human Mint1 MID as bait (residues 222-314). This showed clear binding between the two proteins that was specific for the neuronal Munc18-1 protein as we did not detect any interaction with Munc18-3 (**Fig. 1B**). As the entire N-terminus of Mint1 including the MID is expected to be unstructured in isolation, we hypothesised that Munc18-1 may be binding to a shorter peptide sequence or SLiM (Diella *et al*, 2008) within the MID. To test this, we generated several truncated Mint1 MID sequences and tested their binding to Munc18-1 by GST pull-down assay (**Fig. 1C**). This showed that a twenty-one amino acid region encompassing residues 261-282 was required and sufficient for Munc18-1 binding.

The binding of a synthetic Mint1 peptide to Munc18-1 was next measured by isothermal titration calorimetry (ITC) and the affinity (*K*d) of the interaction was found to be 16.3 ± 4.2 μM (**Fig. 2A**, **Fig. 2B; Table 1**), which is similar to recent reports (Li *et al*., 2023). With a direct assay for Mint1(261-282) peptide interaction we generated a series of N and C-terminal deletions and found that the minimal binding sequence for Munc18-1 in human Mint1 consists of the thirteen residues from Glu267 to Ser280 (**Fig. 2B; Table 1; Fig. S2**). This region is notably the most highly conserved sequence in the N-terminus of Mint1 and Mint2 across species (**Fig. S1**). In line with this we also find that an overlapping peptide from human Mint2 shares a similar binding affinity for Munc18-1 (**Fig. 2B; Table 1; Fig. S2**). Within this region the sequence ^267^EEDIDQIVAE^276^ is invariant across Mint1 and Mint2 homologues from human, fish and xenopus species, although it is not present in worms and flies (**Fig. S1**), and a series of double alanine substitutions in this sequence showed that all these residues are important for Munc18-1 interaction by ITC (**Fig. 2B; Table 1; Fig. S2**). The conservation of this minimal binding sequence suggests that Munc18-1 interactions with Mint1 and Mint2 plays a critical role in the functions of these proteins.

**Figure 2.**
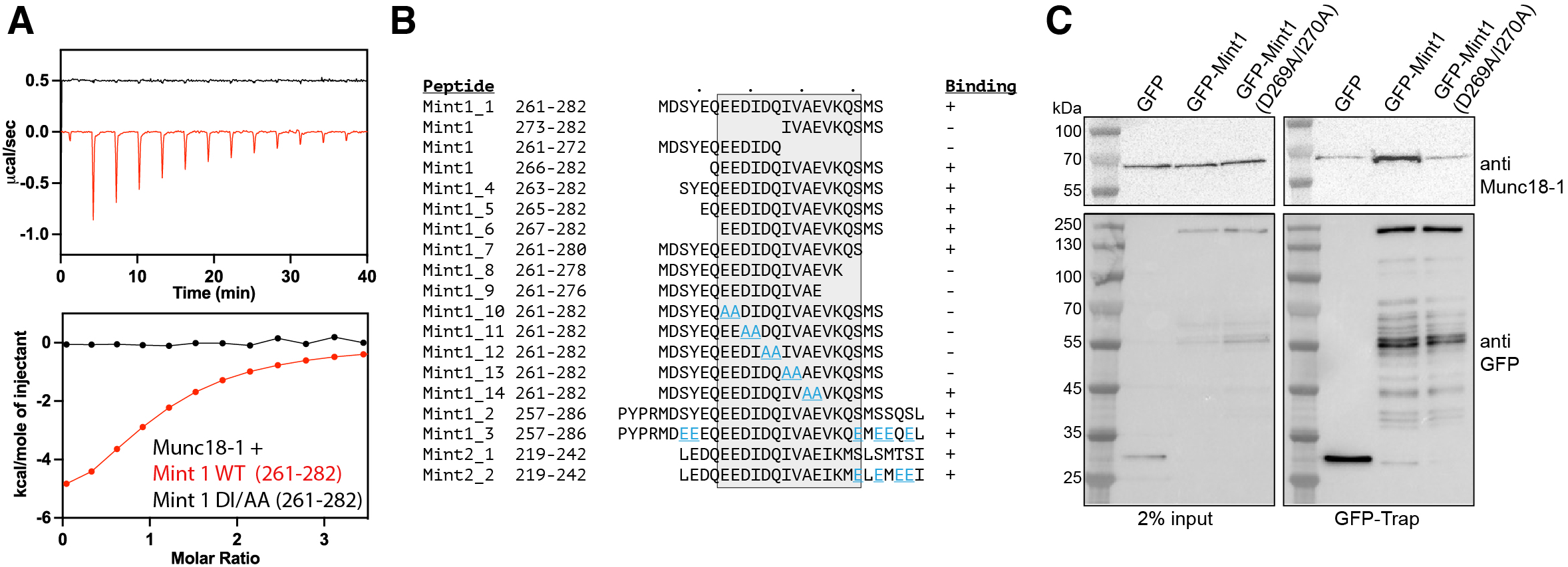
A conserved sequence in Mint1 and Mint2 binds Munc18-1 and is not influenced by phosphorylation. **(A)**ITC of synthetic Mint1^261-282^ peptide binding to purified Munc18-1. The top shows raw ITC data, and the bottom shows integrated and normalised data fit to a 1:1 binding model. **(B)** Sequences of peptides tested for Munc18-1 binding by ITC. The minimal and highly conserved sequence required for Munc18-1 binding is shaded. Mutated peptide residues are highlighted in blue. **(C)** Endogenous Munc18-1 is bound to GFP-tagged Mint1 but not the mutant GFP-Mint1^D269A/I270A^. GFP-tagged Mint1 proteins were transiently transfected into PC12 cells, immunoprecipitated with GFP-nanobody coupled beads, and the bound proteins probed by Western blot with anti-Munc18-1.

**Table 1:**
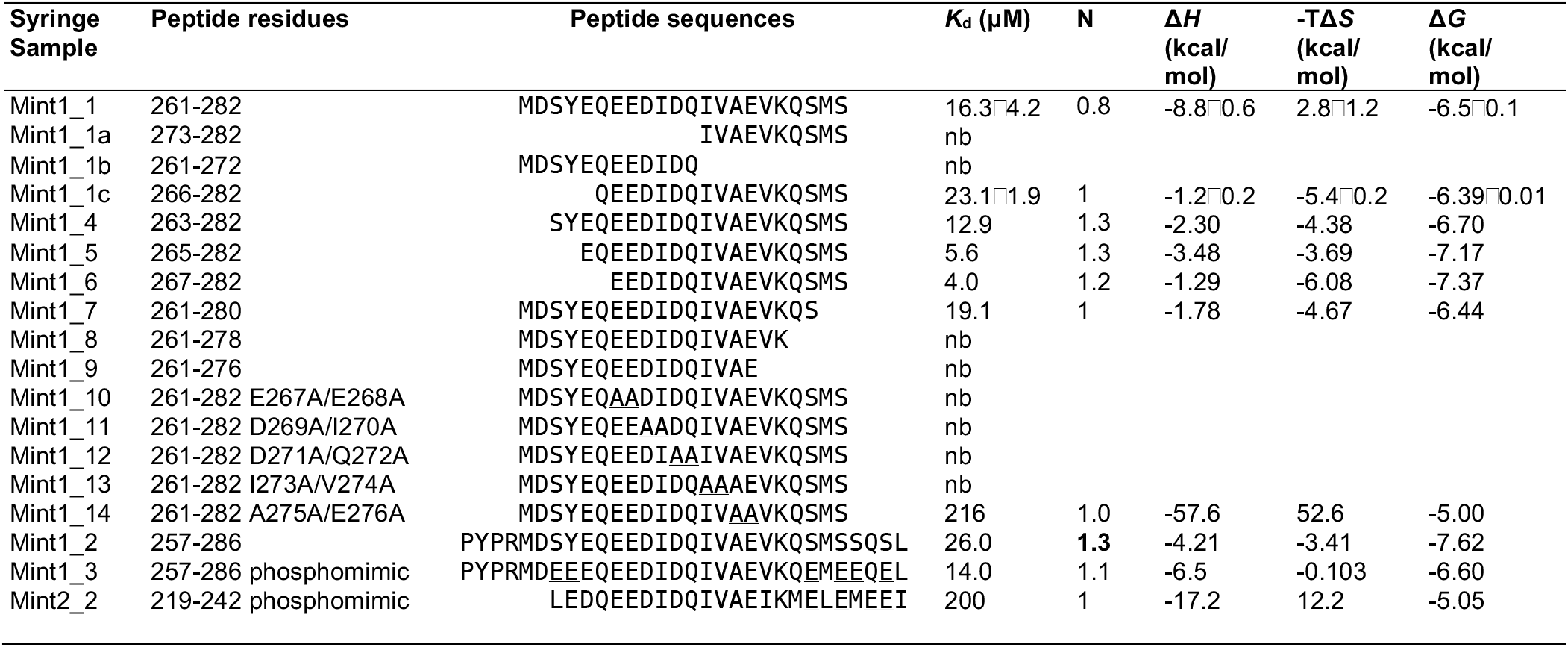
Thermodynamic parameters of Munc18-1 binding to Mint1 by ITC.

We confirmed the binding of Mint1 to endogenous Munc18-1 in PC12 cells by co-immunoprecipitation. GFP-Mint1 or GFP-Mint1(D269A/I270A) were expressed in PC12 cells and bound to GFP-nanotrap beads followed by blotting for the presence of Munc18-1 (**Fig. 2C**). The mutation D269A/I270A reduced Munc18-1 binding in agreement with our *in vitro* ITC data.

### Phosphomimetic Mint1 mutations do not affect the binding to Munc18

The N-terminal region of Mint1 and Mint2 is predicted to be unstructured and can be phosphorylated at numerous sites as catalogued in the PhosphoSitePlus database (Hornbeck *et al*, 2015) (**Fig. S3A**). There are several potential sites of Ser/Thr phosphorylation adjacent to the minimal Munc18-1 binding sequences of Mint1 and Mint2 (**Fig. S3B**). Given the overall negative electrostatic charge distribution of the Mint1 and Mint2 N-terminal domains (**Fig. S3C**), we speculated that adding further negative charges in the form of phosphorylation might influence the affinity of the Munc18-1 interaction. We designed two longer peptides of Mint1 and Mint2 with putative phosphorylated residues altered to phosphomimetic glutamic acid sidechains and tested their binding by ITC (**Fig. 2B; Table 1; Fig. S2**). Our results suggest that phosphorylation of Mint proteins near to the Munc18- interacting sequence does not play a direct role in modulating the Munc18-1 binding affinity at least *in vitro*, although we cannot rule out that Glu is an imperfect mimic of phosphorylation. Previous studies showed that Mint1 and Mint2 N-terminal domains can be phosphorylated upstream of the Munc18-1-interacting sequence by the tyrosine kinase c-Src (Dunning *et al*, 2016). While this enhanced the trafficking of APP via binding the PTB domain, presumably by affecting the overall conformation of Mint1, it had no effect on Munc18-1 interaction. Phosphorylation of Ser236 and Ser238 in Mint2 also enhance the APP interaction (Sakuma *et al*., 2009). These sites lie directly adjacent to the Munc18-1-interacting sequence, but our ITC experiments indicate they do not affect Munc18-1 binding (**Fig. 2B; Table 1; Fig. S2**). Therefore, it appears that while phosphorylation-dependent regulation of the Mint1 N-terminal region plays a role in the interactions with APP (and likely other PTB domain-binding transmembrane proteins such as APLP1, APLP2, Megalin and LRP) they are dispensable for Munc18-1 association *in vitro*.

### Mint1 forms an acidic α-helical motif that binds to Munc18-1 domain 3b

The machine-learning structure-prediction algorithm AlphaFold2 (Evans *et al*, 2022; Jumper *et al*, 2021) has been successfully used to predict structures of protein-peptide complexes by ourselves and others (Ko & Lee, 2021; Poetz *et al*, 2021; Simonetti *et al*, 2021; Tsaban *et al*, 2021). We began our analyses of Mint interactions with a series of modelling experiments to map the complex between Munc18-1 and Mint1 using the ColabFold implementation of AlphaFold2 (Mirdita *et al*, 2022). We initially predicted the complex between the Mint1 and Munc18-1 full-length proteins (**Fig. S4A**). Across multiple models we assessed (i) the prediction confidence measures (pLDDT and interfacial pTM scores), (ii) the plots of the predicted alignment errors (PAE) and (iii) backbone alignments of the final structures. This identified a high-confidence binding sequence in the N-terminal region of Mint1 that correlated precisely with the binding motif identified in our biochemical experiments (**Fig. S4A**).

Based on this initial model and our biochemical mapping of the minimal Mint1 sequence for Munc18-1 binding we performed multiple independent predictions using a shorter peptide region, combined with AMBER energy minimization to optimize amino acid stereochemistry. We found that the minimal sequence of Mint1 was invariably predicted to form an extended α-helical structure that associated with the Munc18-1 domain 3b (**Fig. 3A; Fig. 3B**). This novel binding site is highly conserved in Munc18-1, to a similar degree to the binding site for the Sx1a SNARE protein (**Fig. 3C**). In addition, we found that both the human Munc18-1 and Mint2 proteins and the zebrafish Munc18- 1 and Mint1 orthologues were consistently predicted to form identical structures (**Fig. S4B, S4C**). In contrast, the human Munc18-3 protein was not predicted to form a stable complex with Mint1 (**Fig. S4D**) consistent with their lack of interaction *in vitro* (**Fig. 1B**). Finally, we modelled the tripartite interaction of human Munc18-1 and CASK with an extended Mint1 sequence containing both the Munc18 and CASK-interacting domain (CID) (**Fig. S4E**). The two Mint1 regions were modelled by AlphaFold2 to bind their respective partners in the expected conformations, with the CASK-binding sequence matching closely in structure to the previous crystal structures of the CASK-Mint complex (Wu *et al*., 2020; Zhang *et al*., 2020).

**Figure 3.**
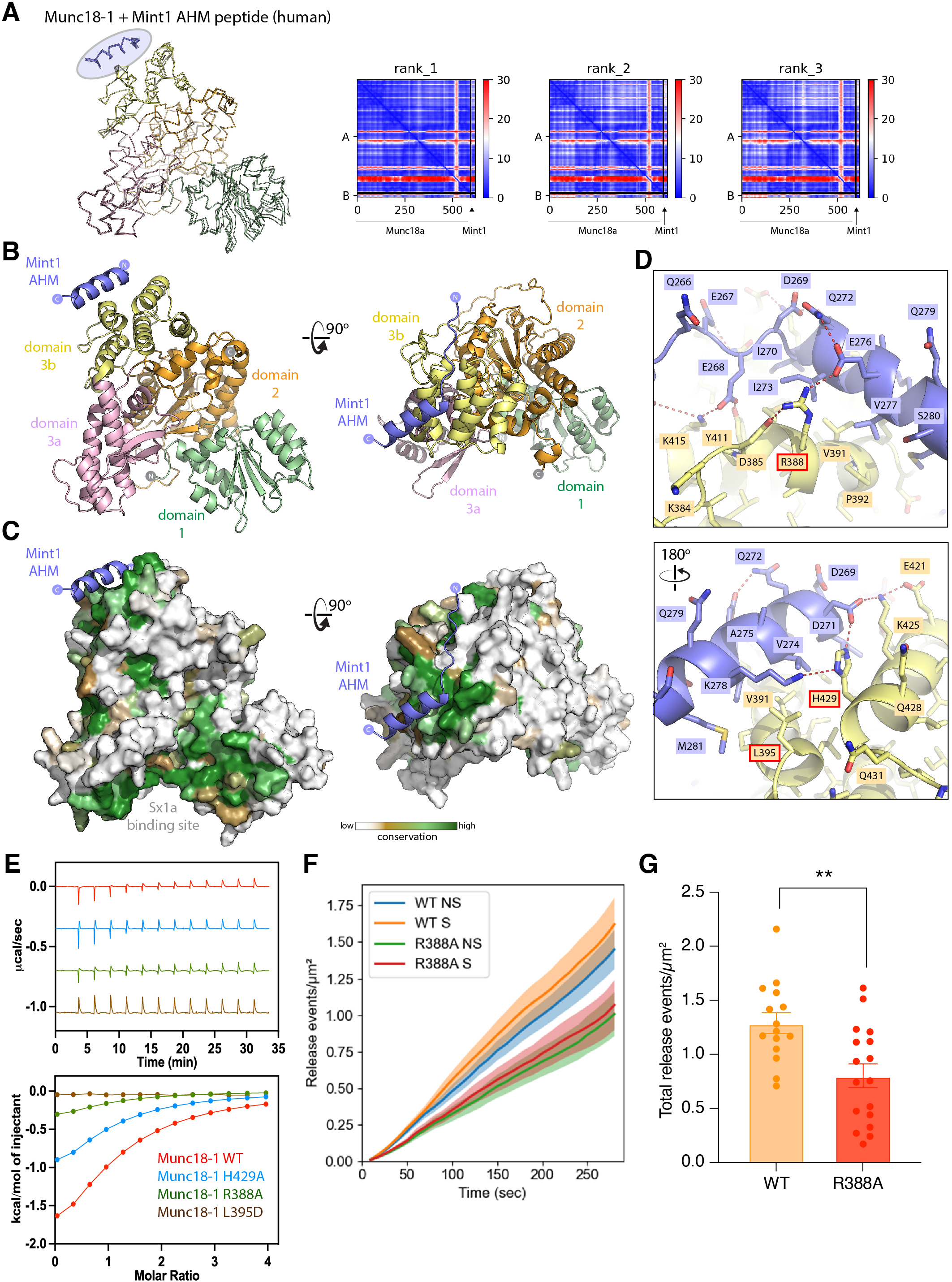
Modelling of Munc18-1 in complex with the Mint1 AHM sequence. **(A)**AlphaFold2 prediction of the complex between Munc18-1 and Mint1 AHM. The three top ranked models are overlaid and shown in backbone ribbon representation. The AHM is consistently modelled in an α-helical structure associated with the Munc18-1 domain3b (highlighted in blue). On the right the predicted alignment error (PAE) is plotted for each model. Signals in the off-diagonal regions indicate strong structural correlations between residues in the peptide with the Munc18-1 protein. **Fig. S4** shows predictions of the full-length Munc18-1 and Mint1 complex as well as models of other Munc and Mint homologues and orthologues. (**B**) The top-ranked complex of Munc18-1 and the Mint1 AHM is shown in cartoon representation, with Munc18-1 domains highlighted. (**C**) As in (**B**) but the surface of Munc18-1 is shown coloured for sequence conservation as calculated by Consurf (Ashkenazy *et al*., 2016). (**D**) Opposite views showing details of the Mint1 AHM bound to the Munc18-1 domain3b. (**E**) ITC of synthetic Mint1^261-282^ peptide binding to purified Munc18-1 and structure-based mutants. The top shows raw ITC data, and the bottom shows integrated and normalised data fit to a 1:1 binding model. Mutated residues are highlighted in panel (**D**). (**F**) Cumulative release events over time for each data group was analysed and the number of release events at each 10 s interval from 0-290 s was determined, plotted as Mean±SEM. NS, non-stimulated; S, stimulated with 2 mM BaCl2. (**G**) Total evoked release events following stimulation were measured per μm^2^. Non-parametric Mann Whitney U test, p < 0.05, *, p <0.01. N = 15 cells (WT) and 17 (R388A) from independent experiments.

The Mint1 interacting sequence forms what we refer to as an acidic α-helical motif (AHM) as proposed by Li and colleagues (Li *et al*., 2023) (see below), and makes a number of critical contacts with Munc18-1 domain3b including both hydrophobic and electrostatic interactions between conserved sidechains (**Fig. 3D; Movie S1**). Consistent with our truncation and mutation analyses of the Mint1 MID peptides (**Fig. 2B**), all predicted core contacts with Munc18-1 are mediated by Mint1 AHM residues Glu262-Ser280. Towards the N-terminus of Mint1 acidic Glu267, Glu268, Asp269 and Glu276 each form complementary bonds with Munc18-1 residues, most notably with Arg388, Tyr411, and Lys415. These are supported by buried hydrophobic interactions of Mint1 Ile270 and Ile273. On the opposite side of the Mint1 α-helix C-terminus a network of bonds is formed between Mint1 Asp271 and Lys278, with Glu421, Lys425 and His429 of Munc18-1. To confirm the predicted binding site of Mint1 and Mint2 we mutated several residues in domain 3b of Munc18-1 including R388A, L395D, and H429A. In ITC experiments all three mutations showed a reduction in binding affinity and enthalpy, with L395D showing an almost complete loss of association (**Fig. 3E**). Altogether the Mint1 AHM is predicted to form an extensive complementary interface with Munc18-1, and we speculate that the relatively modest affinity between the two proteins may in part be due to the entropic cost of the induced α-helical folding of the AHM sequence.

To test the functional importance of this interaction for Munc18-1 dependent exocytosis in neurosecretory cells, we performed an exocytic release assay using Munc18-1/2 double knockout (DKO) PC12 cell line in rescue conditions (Jiang *et al*, 2023). We measured exocytic events by total internal reflection fluorescence (TIRF) microscopy in DKO-PC12 cells co-transfected with VAMP2- pHluorin and either Munc18-1^WT^-mEos3.2 or Munc18-1^R388A^-mEos3.2 – a mutation blocking Mint-1 interaction. VAMP2-pHluorin is classically used to assess vesicular fusion as the intraluminal pH-sensitive pHluorin moiety undergoes unquenching upon exposure to the neutral extracellular environment. This unquenching can be used to study vesicle fusion events and assess the contribution of Munc18-1/Mint1 binding to exocytosis. To assess potential fusion events, we developed a custom Python pipeline that detected puncta of fluorescently labelled vesicles and assessed them over time. A representative cell shows the initiation and disappearance of several vesicles indicative of fusion events in 3D (time being the 3^rd^ axis) (**Fig. S5**). We found that the cells expressing the Mint1 binding deficient Munc18-1^R388A^-mEos3.2 showed a reduced number of exocytic events (**Fig. 3F**) and yielded an approximate 40% decrease in the number of evoked exocytic events relative to the level found upon re-expression of the wild-type protein (**Fig. 3G**).

### The AlphaFold2-predicted interaction precisely matches the crystal structure of the Mint1-Munc18-1 complex

Despite attempts to crystallise Munc18-1 bound to various Mint1 peptides we were unable determine a high-resolution structure of this complex, including with stabilized Munc18-1 variants (Peleg *et al*, 2021). However, as this work was being completed the Song and Feng labs published similar findings regarding the interaction of Munc18-1 with Mint proteins (Li *et al*., 2023). The crystal structure of rat Mint1(227-303) bound to a complex of Munc18-1 and Sx1a was resolved at 3.2 Å resolution, with electron density observed for Mint1 residues 266-283 associated with Munc18-1 domain 3b. This experimental structure correlates precisely with the region of human Mint1 we have mapped biochemically by truncations and mutagenesis and structurally with AlphaFold2. Based on their crystal structure, Li and colleagues termed the Munc18-1-binding Mint1 sequence the acidic α- helical motif (AHM) and we have also adopted this terminology. Overlay of the Munc18-1/Mint1 crystal structure with the top-ranked AlphaFold2 model shows an essentially identical binding mode in all key details (**Fig. 4A**), and it is important to note the crystal structure was not included in the AlphaFold2 training set. One minor difference is that the AlphaFold2 predictions consistently model stable electrostatic contacts involving Mint1 Glu267 and Glu268. These are not seen in the crystal structure, and this is likely because these electrostatic interactions are relatively transient and thus not observed in the modest resolution electron density maps.

**Figure 4.**
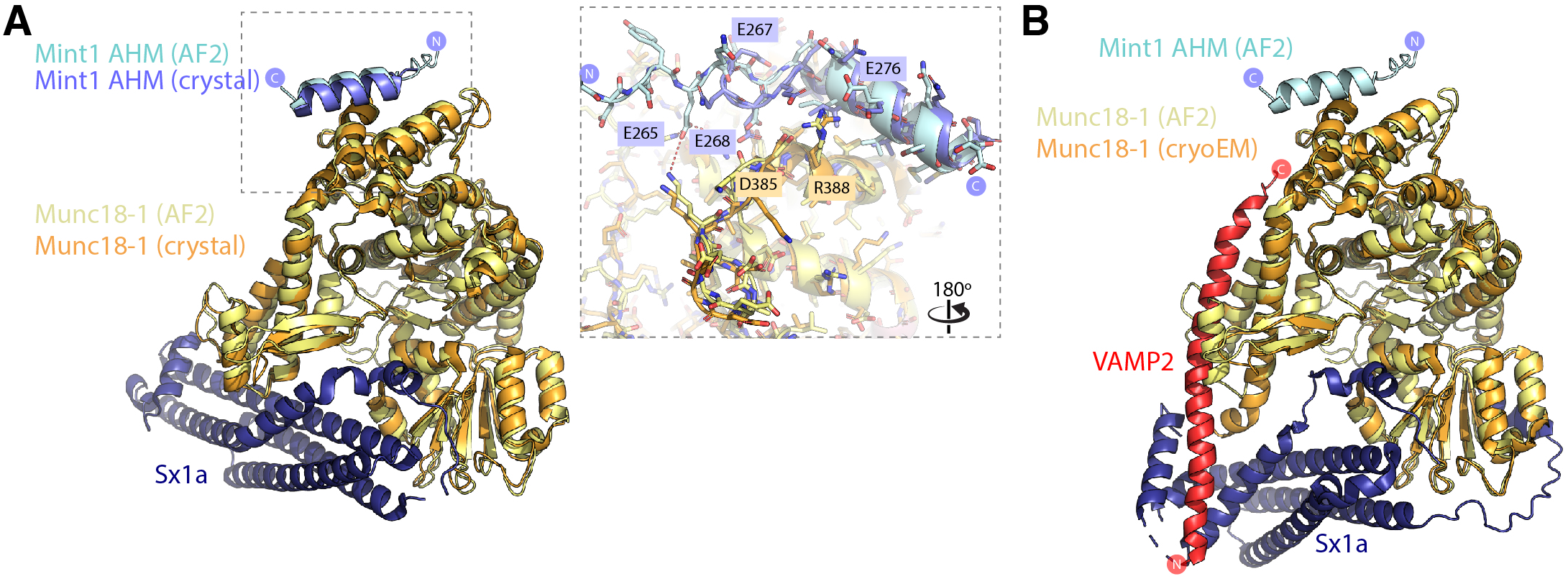
Comparison of the predicted Munc18-1/Mint1 complex with experimental structures. (**A**) Overlay of the Munc18-1 complex with Mint1 AHM predicted by AlphaFold2 and the recent crystal structure of the Munc18-1/Sx1a/Mint1 complex (Li *et al*., 2023) (PDB:7XSJ). The inset shows details of the binding site modelled by AlphaFold2 and observed in the crystal structure. The two structures are identical in all key respects. (**B**) Overlay of the Munc18-1 complex with Mint1 AHM predicted by AlphaFold2 and the cryoEM structure of the Munc18-1/Sx1a/VAMP2 complex (Stepien *et al*., 2022). The Mint1 AHM is expected to bind Munc18-1 independently of the Sxa1 t-SNARE and VAMP2 v- SNARE proteins.

In addition to the crystal structure of Munc18-1/Sx1a bound to the Mint1 AHM, the structure of Munc18-1 was recently determined in a ternary complex with Sx1a and the vesicular R-SNARE VAMP2 (also known as synaptobrevin) by cryoEM (Stepien *et al*, 2022). Similar to what was observed for yeast SM-family protein Vps33, this showed that Munc18-1 can provide a platform to template the assembly of the Qabc-SNARE/R-SNARE complex required for membrane fusion (Baker *et al*, 2015). Overlay of the complexes shows that the Mint1 and VAMP2 binding sites do not overlap and thus Mint1 could potentially associate simultaneously with both SNARE proteins (**Fig. 4B**).

### Mint1 binding to Munc18-1 allosterically modulates Sx1a interaction

Although the Mint-binding site on Munc18-1 does not overlap with either the known VAMP2 or Sx1a binding sites, it is still possible that protein dynamics or allosteric effects could be involved in Mint interaction. To partially address this question, we examined the impact of domain3a deletions and/or the presence of Sx1a on their binding. Previous studies have shown that the flexible hinge-loop region of Munc18-1 domain3a (residues 317-333) is required for efficient priming of secretory vesicles and controls the mobility of Sx1a and subsequent assembly of the SNARE complex (Han *et al*, 2013; Kasula *et al*, 2016; Martin *et al*, 2013). Structural studies of Munc18-1 show that this hinge-loop adopts a ‘closed’ or inhibitory conformation when Munc18-1 is bound to the Sx1a Habc and SNARE domains (Burkhardt *et al*, 2008; Burkhardt *et al*, 2011). Other structures of apo squid Munc18-1, rat Munc18-1 bound to a short Sx1a N-terminal peptide, and the recent cryoEM structure of Munc18-1 in ternary complex with Qa-SNARE Sx1a and R-SNARE VAMP2 show that domain3a can also adopt an ‘open’ conformation that is thought to be necessary for both releasing Sx1a inhibition and providing a platform for binding and assembly of other SNAREs (Bracher *et al*, 2000; Bracher & Weissenhorn, 2001; Hu *et al*, 2011; Stepien *et al*., 2022). This is similar to what is seen when yeast SM protein Vps33 is bound to the Nyv1 SNARE (Baker *et al*., 2015). Surprisingly, we find that deletion of the hinge-loop region in Munc18^Δ317-333^, which has only a modest effect on Sx1a binding (Han *et al*., 2013; Martin *et al*., 2013), abolishes the binding of Mint1 both in GST pull-down and ITC experiments (**Fig. 5A**, **Fig. 5B**).

**Figure 5.**
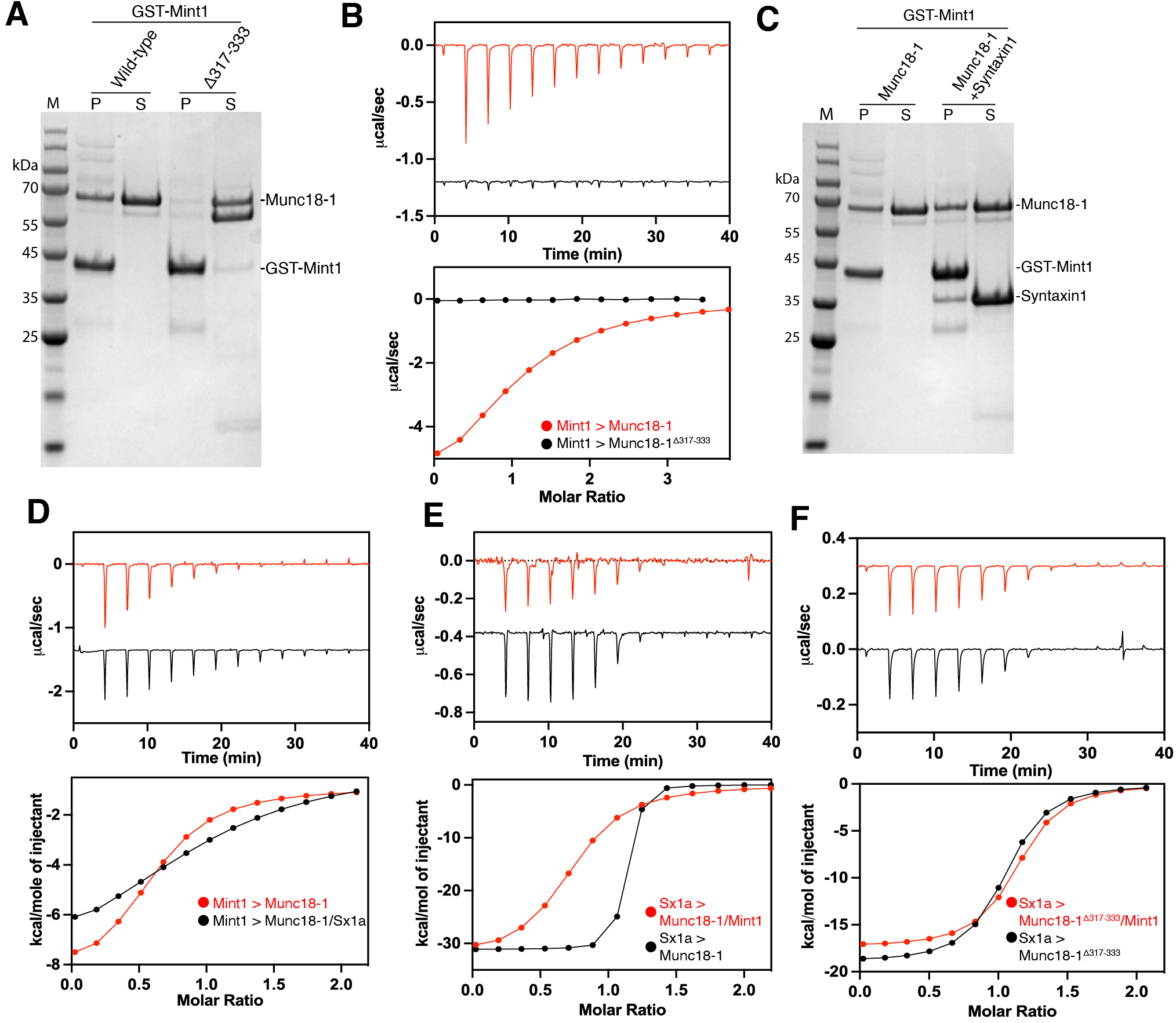
Mint1 and Sx1a show allosteric effects on binding to Munc18-1. **(A)**Pulldowns with GST-Mint1 MID show that Munc18-1 domain3a hinge loop is important for binding. As Mint1 does not contact domain3a this suggests an allosteric effect on the domain3b binding site. Image shows Coomassie stained gel. **(B)** ITC of synthetic Mint1^261-282^ peptide binding to purified Munc18-1 (red) and Munc18-1^β317-333^ (black) confirm the requirement of domain3a for Mint1 interaction. **(C)** Pulldowns with GST-Mint1 MID show that Mint1 can bind Munc18-1 both alone and in the presence of Sx1a. Image shows Coomassie stained gel. **(D)** Although Mint1 and Sx1a can bind Munc18 simultaneously, ITC of Mint1 Mint1^261-282^ AHM peptide binding to Munc18-1 in the absence (red) and presence (black) of Sx1a shows a reduction in binding affinity and enthalpy. **(E)** ITC of Sx1a binding to Munc18-1 in the absence (black) and presence (red) of synthetic Mint1^261-282^ peptide. Together this shows there is a subtle allosteric inhibition of Sxa1 binding to Munc18-1 in the presence of Mint1. **(F)** ITC of Sx1a binding to Munc18-1^β317-333^ in the absence (black) and presence (red) of synthetic Mint1^261-282^ peptide.

As Mint1 binds to Munc18-1 domain 3b, we hypothesized that the perturbed Mint1 interaction on deletion of the distal domain 3a hinge-loop might be due to altered structural dynamics in the combined domain3a/3b module. We therefore tested the binding of Sx1a to Munc18-1 in the absence and presence of the Mint1^261-282^ peptide to determine if there were any changes in Sx1a affinity due to allosteric interactions. By GST pulldown of GST-Mint1 MID we did not observe a gross impact on the ability to bind Munc18-1 in the presence of the high affinity Sx1a ligand (**Fig. 5C**). This is in line with the ability to co-crystallise the three proteins when excess Mint1 peptide is present (Li *et al*., 2023). However, when we quantified the binding affinity by ITC in the presence of Sx1a we saw a small but reproducible reduction in the affinity and enthalpy of binding of Mint1^261-282^ peptide (**Fig. 5D; Table 2**). In reverse experiments, in the presence of a molar excess of Mint1^261-282^ peptide we observe a reciprocal reduction in Sx1a binding affinity (*K*d) from 7.9 nM to 269 nM (**Fig. 5E; Table 2**). This reduced affinity for Sx1a caused by Mint1-dependent allostery is not seen when we use Mint-binding deficient Munc18^Δ317-333^ as expected (**Fig. 5F; Table 2**). Overall, the data indicates that Sx1a and Mint1 binding to Munc18-1 domains 3a and 3b respectively can allosterically regulate the interaction of the other protein. This has potential implications for a role of Mint proteins in Munc18- 1 mediated SNARE assembly, which is a tightly regulated and highly dynamic process.

**Table 2:** Thermodynamic parameters of Munc18-1 binding to Sx1a and Mint1(261- 282) by ITC.

| Syringe sample | Cell Sample | $K_d$ (μM) | N | $\Delta H$<br>(kcal/mol) | $-T\Delta S$<br>(kcal/mol) | $\Delta G$<br>(kcal/mol) | Fold<br>increase in<br>$K_d$ |
| --- | --- | --- | --- | --- | --- | --- | --- |
| Sx1a | Munc18-1 | 0.07±0.2 | 1 | -22.5±2.4 | 11.4±2.3 | -11.1±0 | 1 |
| Sx1a +Mint1 | Munc18-1 +Mint1 | 0.27±0.5 | 1 | -29.7±1.9 | 20.7±2.4 | -9.0±0.3 | 35 |
| Mint1 | Munc18-1 | 8.7±0.2 | 1 | -7.9±0.01 | 0.9±0.02 | -6.9±0.01 | 1 |
| Mint1 | Munc18-1 +Sx1a | 20.6±0.7 | 1 | -8.9±1.9 | 2.5±1.7 | -6.4±0.1 | 2 |
| Sx1a | Munc18-1Δ317-333 | 0.11 | 1 | -18.6 | 9.1 | -9.5 | 1 |
| Sx1a +Mint1 | Munc18-1Δ317-333<br>+Mint1 | 0.09 | 1 | -18.8 | 9.2 | -9.6 | 1 |

### Assessing the network of Mint interactions using AlphaFold2 based predictions

In parallel to the successful modelling of the interaction with Munc18-1/2, we also explored the potential of AlphaFold2 to screen for, and map the binding sites of, other protein-protein interactors of the Mint1 and Mint2 neuronal proteins. Putative Mint1 and Mint2 interactors from the BioGRID repository (Oughtred *et al*, 2021) were screened for direct complex formation with Mint1 and Mint2 respectively using the ColabFold Batch implementation of AlphaFold2 (Mirdita *et al*., 2022) (**Table S1; Table S2**). To assign a direct ‘interactor’ from these *in silico* analyses, we used an approach similar to recent work by Sifri and colleagues (Sifri *et al*, 2023). We initially assessed both the AlphaFold2-derived interfacial PTM (iPTM) score and the resultant PAE graphs which provide confidence metrics for the interactions between the proteins. For those with promising scores, we also examined the predicted structures in PyMOL to assess whether interacting regions involved the expected complementary hydrophobic, polar, and electrostatic contacts. We initially generated three independent predictions of each putative full-length complex in AlphaFold2 in unsupervised batch mode. We found that a minimum iPTM score of ∼0.3 combined with a strong signal in the PAE plots for inter-molecular structural correlation typically provided a practical indicator of a complex that was suitable for more detailed assessment. In these cases, we subsequently ran at least three modelling experiments focusing on the specific domains of Mint1/Mint2 and the putative interactors that were predicted to interact with each other, to assess whether multiple predictions resulted in physically plausible structures that consistently aligned with each other in PyMOL (**Table S1; Table S2**).

As expected, these *in silico* interaction screens with AlphaFold2 correctly predicted several well characterized interactors, but they also provided validation of a number of prospective binding partners so far only identified in high-throughput screens (**Figure S6; Figure S7; Table S1; Table S2**). Known interactors include the cytoplasmic NPxY motif of APP (Matos *et al*., 2012; Xie *et al*., 2013; Zhang *et al*., 1997), and the CID sequence of CASK (Wu *et al*., 2020; Zhang *et al*., 2020) for which crystal structures have previously been determined. In addition, confident predictions were obtained for several other cytoplasmic sequences of transmembrane proteins previously identified to interact with Mint1 and/or Mint2 including lipoprotein receptors LRP1, LRP2 and LRP8 (Gotthardt *et al*., 2000; He *et al*, 2007), calsyntenin-1 (CSTN1) (Araki *et al*, 2003) and KCNJ12 (Leonoudakis *et al*, 2004). These all utilize variations of the NPxY motif found in APP to associate with the Mint PTB domains (**Fig. 6**). While LRP1, LRP2 and LRP8 each possess canonical NPxY sequences, CSTN1 and KCNJ12 are predicted to bind the same site of the PTB domain through divergent NPME and NELA sequences respectively. One other protein identified in BioGRID was predicted with reasonable confidence to associate directly with Mint1. A putative complex between a C-terminal zinc-finger domain from the large protein WIZ (widely interspaced zinc finger-containing protein) was predicted to form with the tandem PDZ domains of Mint1 (**Fig. S6; Table S1**).

**Figure 6.**
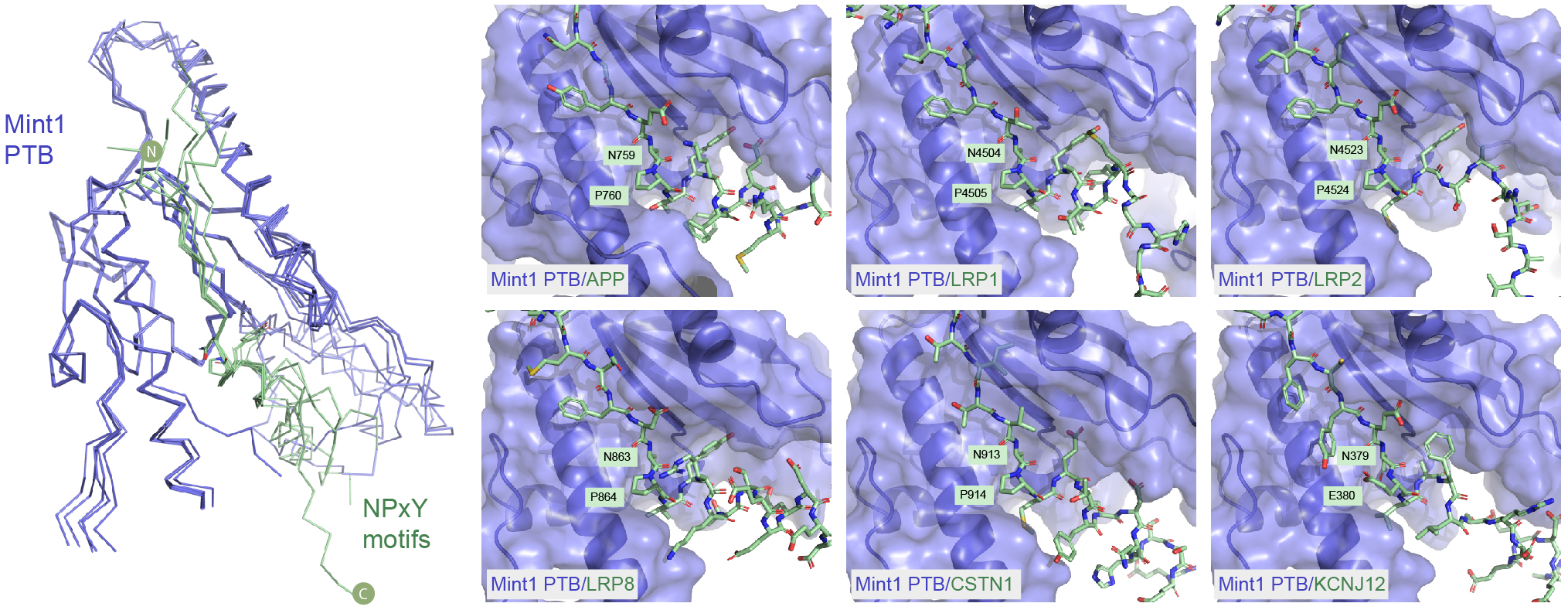
Interactions of the Mint PTB domains with canonical NPxY-containing peptide motifs predicted by AlphaFold2. Overlay of the top-ranked AlphaFold2 predicted structures of the Mint1 PTB domain (blue) in complex with various NPxY-related peptide motifs (green) shown in backbone ribbon representation. These sequences are predicted to bind the canonical binding groove of the PTB domain (with similar interactions predicted for Mint2 (not shown)). The right panel shows details of the different sequences derived from various Mint-interacting transmembrane proteins.

We chose five predicted interactions to describe in more detail; (i) the association of the Mint1 and Mint2 PTB domains with a peptide sequence from the coiled-coil protein Tight Junction Associated Protein 1 (TJAP1) also called Protein incorporated later into tight junctions (PILT) or tight junction protein 4 (TJP4)) (**Fig 7**), (ii) the association of the Mint1 and Mint2 tandem PDZ domains with ADP-ribosylation factor (ARF) GTPases ARF3 and ARF4 (**Fig. 8A**), (iii) the interactions of the C-terminal sequence of the neurexin-1 (NRX1) receptor with the PDZ2 domain of Mint1/Mint2 (**Fig. 8B**), (iv) the interaction of an N-terminal YxxΦ motif in Mint1 with the μ3 subunit of the AP3 clathrin adaptor complex (**Fig. 8C**), and lastly the interaction of a Tyr-containing sequence in the N-terminus of Mint2 with the neuronal ARF GTPase activating proteins (GAPs) IQSEC1 and IQSEC2 (**Fig. 8D**).

**Figure 7.**
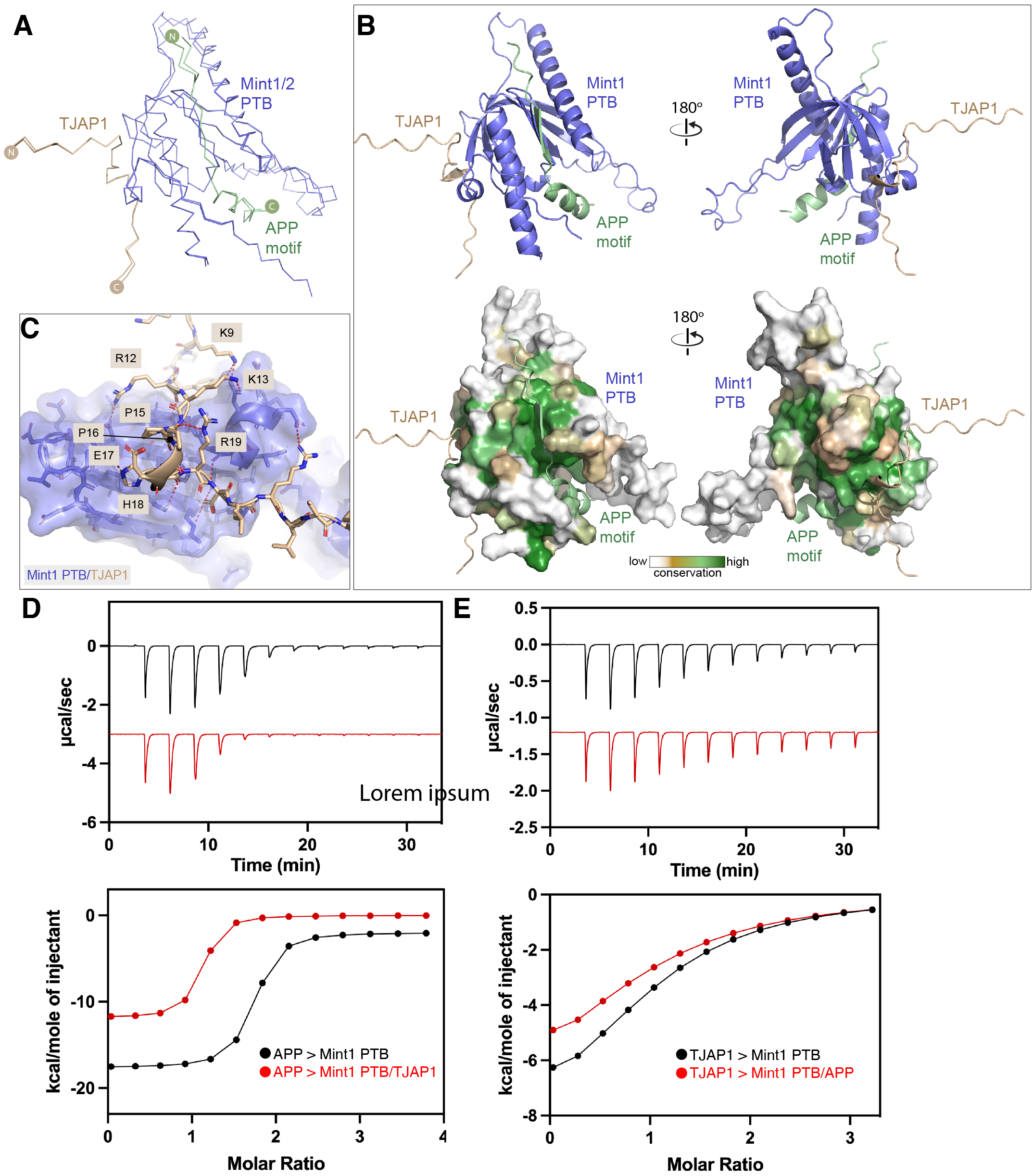
Interactions of the Mint PTB domains with a non-canonical peptide motif from TJAP1 predicted by AlphaFold2. (**A**) Overlay of the top-ranked AlphaFold2 predicted structures of the Mint1 and Mint2 PTB domains (blue) in complex with the APP NPxY motif (green) and the N-terminal peptide of TJAP1 (brown) shown in backbone ribbon representation. (**B**) The top-ranked complex of Mint1 PTB domain bound to APP and TJAP1 is shown in cartoon representation. The lower panel shows the surface of Mint1 coloured for sequence conservation as calculated by Consurf (Ashkenazy *et al*., 2016). (**C**) Details of the Mint1 PTB domain interaction with the TJAP1 peptide. (**D**) ITC of Mint1 PTB binding to the peptide motif from APP in the presence and absence of a peptide from TJAP1. (**E**) ITC of Mint1 PTB binding to the peptide motif from TJAP1 in the presence and absence of a peptide from APP. The top shows raw ITC data, and the bottom shows integrated and normalised data fit to a 1:1 binding model.

**Figure 8.**
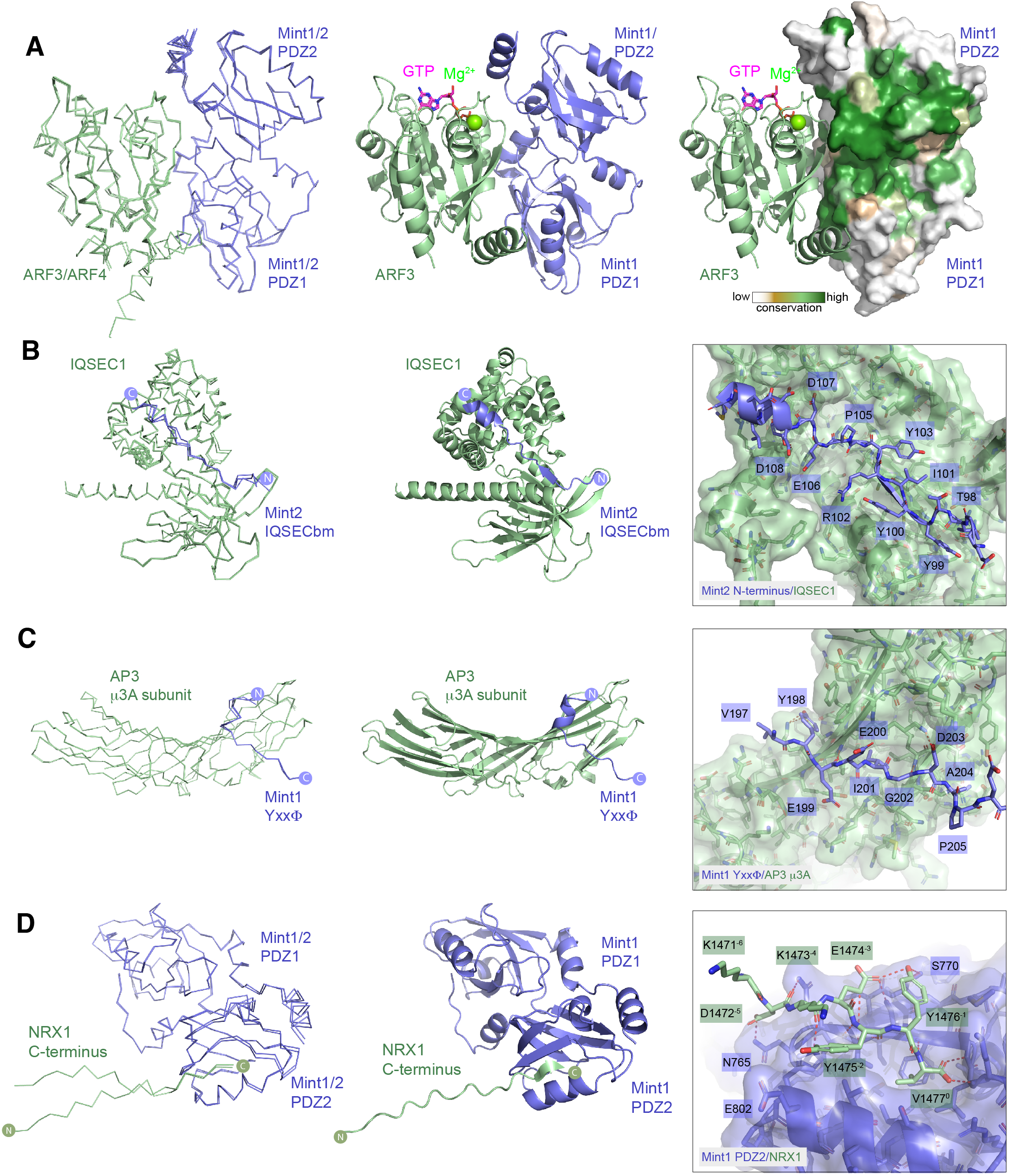
Interactions of the Mint N-terminus and PDZ domains with novel binders predicted by AlphaFold2. (**A**) Overlay of the top-ranked AlphaFold2 predicted structures of the Mint1 and Mint2 tandem PDZ domains (blue) in complex with ARF3 and ARF4 (green) shown in backbone ribbon representation. The middle panel shows the Mint1 complex with ARF3 in ribbon representation, with the position of GTP and Mg^2+^ based on the previous crystal structure of active ARF3-GTP (Lee *et al*., 2019). The right panel shows the same image but with Mint1 surface coloured for sequence conservation as calculated by Consurf (Ashkenazy *et al*., 2016). (**B**) Overlay of the top-ranked AlphaFold2 predicted structures of the N-terminal Mint2 IQSEC binding motif (IQSECbm) (blue) in complex with the C- terminal Sec7 and PH domains of IQSEC1 (green) shown in backbone ribbon representation. The middle panel shows the top ranked Mint2 complex with IQSEC1 in ribbon representation. The right panel inset shows the details of the Mint2 interaction with IQSEC1 PH domain. (**C**) Overlay of the top three ranked AlphaFold2 predicted structures of the Mint1 Yxx<λ motif (blue) in complex with the C-terminal MHD of the AP3 μ3A subunit (green) shown in backbone ribbon representation. The middle panel shows the Mint1 complex with μ3A in ribbon representation. The right panel inset shows the details of the Mint1 Yxx<λ motif interaction with μ3A. (**D**) Overlay of the top-ranked AlphaFold2 predicted structures of the Mint1 and Mint2 tandem PDZ domains (blue) in complex with the C- terminal PDZbm of NRX1 (green) shown in backbone ribbon representation. The middle panel shows the Mint1 complex with ARF3 in ribbon representation. The right panel inset shows the details of the NRX1 interaction with Mint1 PDZ2 domain.

Outside of the canonical interactions of NPxY-related motifs with the Mint1 and Mint2 PTB domains, we were intrigued by a high confidence prediction involving a short peptide sequence in TJAP1 (previously reported in high-throughput proteomic screens (Huttlin *et al*, 2021) (**Fig. S7; Table S2**). The TJAP1 binding site is distinct from the binding groove of NPxY motifs, and subsequent predictions of the Mint1 and Mint2 PTB domains in the presence of both the NPxY motif of APP, and the peptide sequence from TJAP1 show highly consistent dual peptide interactions on opposite faces of the PTB domain (**Fig. 7A, B**). Both the APP and TJAP1-binding surfaces of Mint1 and Mint2 are highly conserved (**Fig. 7B**). The binding sequence of TJAP1 encompasses N-terminal residues ^9^KPYRKAPPEHRELR^22^, with buried aliphatic sidechains, and complementary electrostatic and hydrogen-bond contacts as shown in **Fig. 7C**. The sequence ^16^PEHR^19^ is predicted to form a β- turn structure where the Pro16 sidechain forms a stacking interaction with the Arg19 guanidino group. To validate these AlphaFold2 predictions we confirmed the binding of TJAP1 to Mint1 experimentally by ITC. Using the NPxY-containing sequence ^750^SKMQQNGYENPTYKFFEQMQ^769^ of APP as a positive control we confirmed that this bound to the Mint1 PTB domain with an affinity (*Kd*) of 0.3 μM (**Fig. 7D; Table 3**), similar to the affinity reported previously (Matos *et al*., 2012). The TJAP1 peptide ^9^KPYRKAPPEHRELR^22^ bound to the Mint1 PTB domain with a modest affinity (*K*d) of 20.9 μM (**Fig. 7E; Table 3)**. Importantly, in competition experiments the presence of the APP peptide did not appreciably alter the TJAP1 affinity nor vice versa, confirming that the two peptides interact with distinct sites on Mint1 as predicted (**Fig. 7D and 7E; Table 3)**. These results show the Mint PTB domains are capable of recruiting proteins via two distinct peptide-motifs.

**Table 3:** Thermodynamic parameters of Mint1 PTB domain binding to TJAP1 and APP peptides by ITC.

| Syringe sample | Cell Sample | $K_d$ (μM) | N | $\Delta H$<br>(kcal/mol) | $-T\Delta S$<br>(kcal/mol) | $\Delta G$<br>(kcal/mol) |
| --- | --- | --- | --- | --- | --- | --- |
| TJAP1(9-22) | Mint1 PTB | 20.9 ± 2.4 | 1 | -8.9 ± 0.9 | 2.5 ± 0.9 | -6.4 ± 0.1 |
| TJAP1(9-22) | Mint1 PTB +<br>APP(750-769) | 29.4 ± 9.5 | 1 | -7.1 ± 1.6 | 1.4 ± 1.8 | -6.2 ± 0.2 |
| APP(750-769) | Mint1 PTB | 0.31 ± 0.01 | 1 | -15.6 ± 0.7 | 6.7 ± 0.6 | -8.9 ± 0.01 |
| APP(750-769) | Mint1 PTB +<br>TJAP1(9-22) | 0.58 ± 0.01 | 1 | -11.8 ± 0.1 | 1.7 ± 1.1 | -8.5 ± 0.01 |

Mint2 was previously identified in a yeast two-hybrid screen for interactors of ARF3 (Hill *et al*, 2003). These screens identified fragments of Mint2 containing the tandem C-terminal PDZ domains, and subsequent experiments showed that Mint1, Mint2 and Mint3 could all interact with both ARF3 and ARF4. AlphaFold2 predictions of full-length Mint2 complexed with ARF3 and ARF4 revealed a very high confidence interaction of the GTPases with the tandem PDZ domains, correlating with the yeast two-hybrid studies (Hill *et al*., 2003) (**Fig. S7; Table S2**). Predictions focused on the Mint1 and Mint2 PDZ domains subsequently produced highly consistent complex structures with the two GTPases (**Fig. 8A**). The structure of activated ARF3-GTP bound to a bacterial toxin called MARTX was previously solved by X-ray crystallography (Lee *et al*, 2019). AlphaFold2 predicts ARF3 and ARF4 to be in the active GTP-Mg^2+^-loaded conformation when bound to Mint1, and like the bacterial effector Mint1 and Mint2 primarily engage the switch 2 and interswitch regions of the GTPases, making little contact with the switch 1 region. The ARF GTPases bind an extensive and conserved surface with the interface composed of regions from both PDZ1 and PDZ2 domains in the tandem PDZ structure.

As well as the ARF GTPases, two neuronal ARF GAPs IQSEC1 and IQSEC2 (IQ motif and SEC7 domain-containing proteins), were also predicted to bind to an N-terminal sequence in Mint2 ^96^GITYYIRYCPEDD^108^ (**Fig. 8B; Fig. S1**). The core binding sequence consists of a several Tyr sidechains from Mint2 that stack along a conserved surface groove in the C-terminal PH domain of IQSEC1 and IQSEC2, with complementary electrostatic contacts also made by downstream Glu and Asp residues from Mint2 with Lys and Arg sidechains in IQSEC proteins. Although the sequence is semi-conserved in Mint1 homologues (**Fig. S1**), AlphaFold2 does not predict a confident interaction with these proteins (not shown), suggesting their sequences are less optimized for binding. The IQSEC1 and IQSEC2 proteins are Ca^+^/CaM-regulated synaptic proteins that can activate all members of the ARF GTPase family (Bai *et al*, 2023; Peurois *et al*, 2017; Um, 2017), and are mutated in neuronal developmental disorders including X-linked intellectual disability with early onset epileptic encephalopathy (Ansar *et al*, 2019; Zerem *et al*, 2016). Notably, the IQSEC1 protein was previously shown to bind a short Tyr-containing sequence in the C-terminus of the GluA2 AMPA receptor via the PH domain (Scholz *et al*, 2010), and we speculate that Mint2 and GluA2 may utilize the same binding site in the IQSEC proteins.

The next predicted interaction we examined in detail was that of Mint1 with the μ3 subunit of the tetrameric AP3 clathrin adaptor complex (**Fig. 8C**), another unexpected association reported in the same high-throughput screens as TJAP1 (Huttlin *et al*., 2021). The μ3 subunit has an N-terminal longin domain that embeds it within the AP3 tetramer, and a C-terminal μ-homology domain (MHD) that associates with YxxΦ tyrosine-based motifs (where Φ is a bulky hydrophobic residue), typically in transmembrane cargos for sorting from endosomes to lysosomal compartments (Mardones *et al*, 2013; Sanger *et al*, 2019). The predicted structure of the Mint1 ^298^YEEI^301^ sequence closely resembles the crystal structure of the TGN38 motif YQRL bound to the rat μ3 MHD, with both Tyr298 and Ile301 inserting into complementary surface pockets as typically seen for YxxΦ motifs. We also tested AlphaFold2 predictions of the Mint1 sequence with other μ-subunits from AP1, AP2, AP4 and AP5. While μ1, μ2 and μ4 proteins were each predicted to form a complex, the iPTM confidence scores were much lower than μ3, and μ5 was not predicted to bind at all.

Although not listed in the BioGRID entries for Mint1 or Mint2, other studies proposed that their PDZ domains interact with C-terminal sequences of NMDA receptors, kalirin-7, Neurexin-1 (NRX1), ApoER2/LRP8 and LDLR (Biederer & Sudhof, 2000; Gotthardt *et al*., 2000; Jones *et al*., 2014; Minami *et al*., 2010; Motodate *et al*., 2019). PDZ domains are small scaffolds that bind to PDZ binding motifs (PDZbms) typically found at the C-terminus of their interacting proteins. They fall in three main classes; Type 1 with consensus [S/T]xΦ (x = any amino acid; <λ = hydrophobic amino acid), Type 2 with consensus <λx<λ, and Type 3 with consensus [E/D]x<λ. (Hung & Sheng, 2002; Nourry *et al*, 2003). We used AlphaFold2 to predict the interaction of the Mint1 and Mint2 PDZ domains with the C-terminal sequence of NRX1, which conforms to a Type 2 PDZbm. This confidently predicted an interaction between the C-terminal ^1471^KDKEYYV^1477^ NRX1 sequence with the second PDZ2 domain of the PDZ1-PDZ2 tandem structure (**Fig. 8D**). The C-terminal Val1477 side chain (position ‘0’ in standard PDZbm nomenclature) docks in a complementary hydrophobic pocket with the terminal carboxyl group hydrogen bonding with the backbone amides of Mint1 Leu759 and Gly760. The NRX1 Tyr1475 and Tyr1476 side chains at positions -1 and -2 pack into complementary surface grooves, while both main-chain and side-hydrogen bonds upstream of the C-terminal interaction provide further specificity. This requires experimental validation, but combined with the fact that the C-terminal Mint1 sequence PVYI (PLIY in Mint2) itself can form an intramolecular *cis-*interaction with its own PDZ1 domain (Long *et al*, 2005) it suggests that the PDZ2 domain of the Mint proteins provides the major platform for recruiting PDZbm-containing *trans-* ligands.

While we have focused on proteins that *are* predicted to bind to Mint1/Mint2, it is notable that from the list of putative BioGRID interactors the majority are *not* predicted to associate directly with the Mint adaptors, including many that have previously been identified using methods such as co-immunoprecipitations (**Table S1; Table S2**). In some case this could be due to limitations with the predictive ability of AlphaFold2, or potentially other requirements such as post-translational modifications not accurately represented in AlphaFold2 predictions. However, propose that many of the proteins reported in BioGRID either bind indirectly (via other proteins not included in our binary predictions) or are non-specific interactions detected by the proteomics methods.

## DISCUSSION

The interaction of Mint proteins with Munc18-1 has long been known to be important for synaptic neurotransmitter and hormonal release through the regulation of SNARE-mediated vesicle fusion. While the molecular basis for the scaffolding and trafficking of the APP transmembrane protein and CASK adaptor have been structurally characterised (Matos *et al*., 2012; Wu *et al*., 2020; Xie *et al*., 2013; Zhang *et al*., 1997; Zhang *et al*., 2020), until recently the mechanism by which Mints bind to Munc18-1 was unknown. In this work, and in a recent study by Li and colleagues (Li *et al*., 2023), the interaction is revealed to be via the binding of Munc18-1 domain3b to a conserved α-helical motif (AHM) in the N-terminal unstructured domains of the Mint1 and Mint2 neuronal homologues. The binding surface on Munc18-1 is distinct from its known binding sites for the Sx1a and VAMP2 SNARE proteins, although we find there is a small but significant reduction in the binding affinity of Sx1a in the presence of the Mint1 AHM. In line with a role for the Mint1 interaction in Munc18-1-dependent exocytosis we observed that perturbation of Munc18-1 Mint-1 interaction in domain 3b impacted exocytosis in neurosecretory cells. We speculate this could point to a role of Mint proteins in regulating the SNARE complex dynamic templating activity of Munc18-1 which will be worth future investigation.

The AHM sequence found in Mint1 and Mint2 is conserved across many species, although it appears not to be present in some organisms such as nematodes and flies. Further, it may be relatively specific to the Mint proteins, with few if any other proteins possessing similar motifs. We scanned the human genome using ScanProsite and did not find any other proteins with highly similar sequences. Li and colleagues reported potential AHMs in Munc13-1, Bassoon, and Atg16L, although no binding was detected using the putative motif from Munc13-1 by ITC (Li *et al*., 2023). Future proteomic studies of Munc18-1 using specific domain3b mutations may confirm the existence of other proteins able to bind this site, but there are unlikely to be a large number. Like the CASK-binding CID sequence in Mint1, the Mint1 and Mint2 AHM sequences lie within their extended and intrinsically unstructured N-terminal domain, likely adopting their α-helical structures via induced folding upon Munc18-1 interaction. This may explain the relatively modest binding affinity for Munc18-1 despite the reasonably large binding surface. The interaction between these proteins is thus likely highly context dependent, relying on both their specific coupling as well as their colocalization at the membrane surface and likely clustering with other proteins such as the SNAREs, APP, neurexins, and potentially small GTPases like ARF3.

In addition to dissecting the mechanism of Mint1 and Mint2 interaction with Munc18-1, we have used machine learning-based structure prediction with AlphaFold2 to assess the broader interactome of the Mint proteins. These predictions provide insights into those interactions that are likely to be *directly* mediated by the Mint proteins, as validated in one instance with the direct binding of TJAP1 confirmed. Our results thus provide an example of how AlphaFold2 and similar algorithms can be used as a type of triage of large proteomic datasets, providing additional confidence in the plausibility of direct interactions (Burke *et al*, 2023; Gao *et al*, 2022a; Gao *et al*, 2022b; Humphreys *et al*, 2021; Sifri *et al*., 2023; Yu *et al*, 2023). Such an approach has the potential to inform and accelerate subsequent experimental validation of molecular complexes detected in high-throughput screens, by providing greater assurance as to which hits represent specific interactors.

Apart from the expected predictions of CASK and APP, for which previous crystal structures are available, there were several other notable complexes that were confidently modelled by AlphaFold2 in this study. A number of proteins have been reported to interact with the PDZ domains of Mint1 and Mint2 via C-terminal PDZbms, including calcium channels (Maximov *et al*, 1999), kalirin-7 (Jones *et al*., 2014; Penzes, 2001), NMDA receptors (Motodate *et al*., 2019) and NRX1 (Biederer & Sudhof, 2000; Dean & Dresbach, 2006). Furthermore, the C-terminus of Mint proteins can form an intramolecular interaction with their own PDZ1 domain, acting as an autoinhibitory sequence of PDZ1 (Long *et al*., 2005). Taking NRX1 as an example, we found that its Type II PDZbm was strongly predicted to interact with the PDZ2 domain in the canonical β-strand orientation, and since other neurexin homologues share the same C-terminal sequence they likely use the same binding mechanism. This hypothesis, as well as whether the second PDZ2 domain is bound by other PDZbms warrants further experimental studies.

One novel Mint interactor predicted by AlphaFold2 and validated in direct binding experiments was TJAP1, which was previously identified in high-throughput proteomic screens with all three Mint homologues (Huttlin *et al*., 2021). TJAP1 has an N-terminal unstructured region, which we predicted to interact with Mint PTB domains, a central-coil region predicted by AlphaFold2 to form a homodimer (not shown), and an extended C-terminal unstructured and Proline-rich domain. The precise function of TJAP1 is essentially unknown, although it was identified in yeast two-hybrid assays to bind the GTPase ARF6 (Tamaki *et al*, 2012) and discs large-2 (Dlg-2/PSD93) (Kawabe *et al*, 2001), a member of the membrane-associated guanylate kinase (MAGUK) protein family that includes CASK (Ye *et al*, 2018). TJAP1 is localised to both the Golgi and tight junctions (Kawabe *et al*., 2001; Tamaki *et al*., 2012), and the putative interaction with the TJAP1 N-terminal peptide sequence is confidently predicted with not only the neuronal Mint1 and Mint2 proteins but also the ubiquitous Mint3 protein (not shown) so it is probable that the association is important in diverse cell types as well as in neurons.

The ARF3 and ARF4 small GTPases were originally identified to bind Mint proteins in yeast two-hybrid screens (Hill *et al*., 2003), however no subsequent studies have examined the mechanism or functional role of these interactions. Our modelling indicates a conserved binding site involving the PDZ1 and PDZ2 tandem domains of the Mint proteins, which supports the original yeast two-hybrid mapping experiments (Hill *et al*., 2003). ARF3 is highly enriched in the brain, and both proteins play a role in maintaining recycling endosome morphology and integrity (Kondo *et al*, 2012; Nakai *et al*, 2013). Interestingly ARF3 mutations have recently been found in patients with neurodevelopmental disorders characterised by brain abnormalities, microcephaly and seizures (Fasano *et al*, 2022; Sakamoto *et al*, 2021). It is tempting to speculate these disorders may overlap with synaptic pathologies caused by mutations in Munc18-1, CASK and other synaptic proteins, thus suggesting a role for ARF3 in the synaptic vesicle trafficking pathways that could in part be mediated through the Mint proteins.

The last predicted interaction we examined was that of Mint1 with the AP3 clathrin adaptor complex. This involves binding of the AP3 μ3A domain with a canonical YxxΦ sequence in the Mint1 N-terminal region (not present in Mint2 or Mint3). This would indicate that the Mint1 N-terminus has at least two functions distinct from the other Mint isoforms; the ability to bind CASK, and the potential to couple Mint1 and bound proteins (such as APP or neurexins for example) into AP3 mediated transport structures. AP3 is primarily found on endosomes where it mediates trafficking to lysosomes and lysosome-related organelles (Sanger *et al*., 2019), and depending on specific subunit isoforms has important roles in axonal transport and synaptic function by regulating the reformation of synaptic vesicles from endosomes derived from bulk synaptic endocytosis (Blumstein *et al*, 2001; Evstratova *et al*, 2014; Li *et al*, 2016; Newell-Litwa *et al*, 2010; Seong *et al*, 2005; Shetty *et al*, 2013; Voglmaier *et al*, 2006). Similar to ARF3, mutations in neuronal AP3 isoforms can lead to neurodevelopmental disorders with some overlapping features with other synaptopathies (Lanoue *et al*., 2019), suggesting a potential functional overlap of Mint1 and AP3 in synaptic integrity (Ammann *et al*, 2016; Assoum *et al*, 2016).

An overall model for Mint1 is shown in **Fig. 9** summarising the known and predicted interactions mediated by this scaffold protein. **Fig. 9A** shows an AlphaFold2 prediction of the full-length protein highlighting binding sequences and structural domains of the protein and underlines the highly extended nature of the N-terminal intrinsically disordered domain containing the Munc18- 1 binding AHM as well as binding motifs for the AP3 adaptor and CASK. **Fig. 9B** is a cartoon summary of the interactions described above and highlights the overall scaffolding function of this protein. From our biophysical and biochemical studies, we speculate that the reduced affinity of Sx1a for Munc18-1 in the presence of the Mint AHM sequence could lead to an enhancement of SNARE assembly mediated by Munc18-1 templating, although this will require more extensive testing. A final caveat to this model is that it does not account for temporal regulation of the various interactions, the impact of posttranslational modifications or the cellular environment where each interaction is likely to occur including the plasma membrane and other organelles such as endosomes and the Golgi.

**Figure 9.**
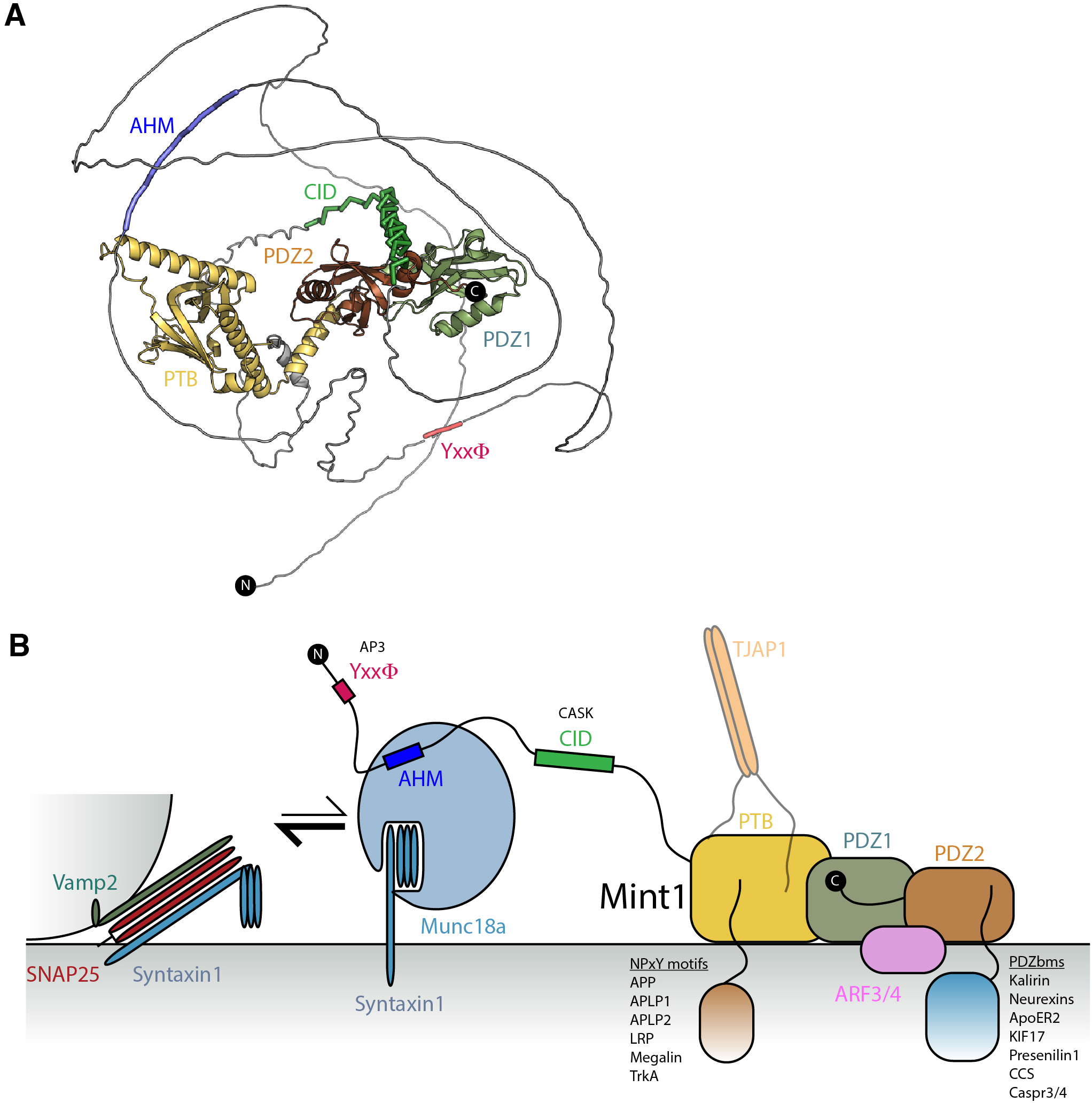
Mint1 structural model and interactions. (**A**) Structural model of Mint1 derived from AlphaFold2 (Ammann *et al*., 2016). (**B**) Schematic summary of Mint1-mediated interactions, and speculative model suggesting that at the cell surface Mint1 may act to reduce the affinity of Munc18-1 for the auto-inhibited Sx1a, thus enhancing the ability of Sx1a to associate with VAMP2 and SNAP25 to form the trans-SNARE assembly required for vesicle fusion. The C-terminal domains of Mint1 in contrast are associated with other proteins containing NPxY and PDZbm sequences and ARF small GTPases that may enhance Mint1 membrane recruitment and modulate transmembrane protein trafficking.

In summary, we have mapped and characterised the specific association of the neuronal Mint1 and Mint2 proteins with the SNARE regulatory protein Munc18-1 providing a high-resolution snapshot for how these key synaptic proteins interact with other, confirming and extending recent related work (Li *et al*., 2023). This study further emphasises the ability of AlphaFold2, at least in many instances, to predict protein-peptide interactions with a high degree of accuracy. By applying a wider set of systematic analyses, our work has revealed likely modes of interaction between the Mint proteins and a variety of known and novel effectors, which provides a foundation for future mechanistic studies of their important role in synaptic activity.

## METHODS

### Antibodies, plasmids and peptides

Human Mint1 sequences for bacterial expression were codon optimised and sub-cloned into the pGEX4T-2 plasmid by Genscript (USA). The constructs generated were GST-tagged Mint1(226- 314)(MID), Mint1(261-272), Mint1(261-282), Mint1(226-289), Mint1(222-314), Mint1(237-289).

Human Mint1 open reading frame and mutant Mint1(D269A/I270A) were obtained from Genscript and cloned into the pcDNA3.1-N-eGFP. The human Mint1 PTB domain (residues 448-623) sequence was synthesised and cloned into pGEX6P-2 by Gene Universal.

All Mint-derived synthetic peptides were purchased from Genscript (USA). The human APP(750-769) (SKMQQNGYENPTYKFFEQMQ) and TJAP1(9-22) (KPYRKAPPEHRELR) peptides were made by solid phase peptide synthesis in-house, purified by reverse phase HPLC and purity assessed by mass spectrometry. GFP polyclonal antibody and Goat anti-mouse secondary antibody were purchased from Thermo Fisher and Mouse monoclonal anti-munc18-1 was purchased from BD biosciences. pmEos3.2-N1 Munc18-1^WT^ was created by restriction digestion of Munc18-1^WT^ (double digestion with NheI/AgeI, New England Biosciences) from pmEos2-N1 Munc18-1^WT^ (Kasula et al. 2015 *Journal of Cell Biology*), vector linearisation of pmEos3.2-N1 with NheI/AgeI was followed by T4 Ligation. pmEos3.2-N1 Munc18-1^WT^ was then used as a template to introduce the following missense mutation R388A using QuickChange Lightning site-directed mutagenesis kit (Agilent Technologies, Cat# 210518) as per manufacturer’s instructions. Primers were designed by PrimerX (site: http://bioinformatics.org/primerx/) and ordered from SigmaAldrich. M18-1_R388A_For: 5’ GAAAAAATCAAGGACCCCATGGCAGCCATTGTCCCCATCCTGC 3’ and M18-1_R388A_Rev: 5’ GCAGGATGGGGACAATGGCTGCCATGGGGTCCTTGATTTTTTC 3’. All new plasmids were verified by Sanger sequencing performed by the Australian Genome Resource Facility (Brisbane, Australia). pCI VAMP2-pHluorin was a kind gift from James Rothman (Miesenbock *et al*, 1998).

### Recombinant protein expression and purification

All proteins were expressed in *Escherichia coli* Rosetta™ BL21(DE3) cells. GST-Mint1 (226-314, the Munc18-1 interacting domain (MID)), containing pGEX4T-2 vector was transformed into Rosetta cells and plated on a LB/Agar plate supplemented with Ampicillin (0.1mg/mL). Single colony was then used to inoculate 50 mL of LB medium containing Ampicillin (0.1 mg/mL). and the culture was grown overnight at 37 ^ο^C with shaking at 180 rpm. The following day, 1L of LB medium containing antibiotics Ampicillin (0.1 mg/mL) and Chloramphenicol (0.034 mg/mL) was inoculated using 10 mL of the overnight culture. Cells were then grown at 37 ^ο^C with shaking at 180 rpm to an optical density of 0.7-0.8 at 600 nm and the protein expression was induced by adding 0.5 mM IPTG (isopropyl-β- D-thiogalactopyranoside). Expression cultures were incubated at 20 ^ο^C overnight. And the cells were harvested next day by centrifugation at 4000 rpm for 15 min using Beckman rotor JLA 8.100. Cell pellets were resuspended in 20 mL (for cell pellet from 1L) of lysis buffer (50 mM Tris, pH 8.0, 500 mM NaCl, 10% glycerol, 1 mM DTT, Benzamidine (0.1mg/mL), and Dnase (0.1 mg/mL)).

Resuspended cells were further lysed by using the cell disrupter (Constant systems, LTD, UK, TS-Series) and the soluble fraction containing GST-Mint1 was separated from cell debris by centrifugation at 18,000 rpm for 30 min at 4 ^ο^C. The soluble fraction was first purified by affinity chromatography using glutathione-Sepharose resin and the GST-Mint1 was eluted using 50 mM Tris, pH 8.0, 500 mM NaCl, 10% glycerol, 1 mM DTT buffer and the protein containing fractions were concentrated and further purified by gel filtration chromatography (Superdex 75 (16/600), GE Healthcare). GST-Mint1 containing fractions were analysed by SDS PAGE. All the other GST-Mint1 N-terminal constructs (GST-Mint1 261-272, 261-282, 226-289, 222-314, 237-289) were also expressed and purified as described above. The GST-Mint1 PTB domain was expressed as above, but the GST tag was removed by incubation with Prescission protease, followed by gel filtration into 50 mM HEPES, 200 mM NaCl, 0.5 mM TCEP and 5% glycerol.

For GFP-nanotrap preparation, the plasmid pOPINE harbouring His-SUMO-GFP-nanotrap was transformed into *Escherichia coli* Bl21 (DE3) cells and plated on a LB/Agar plate supplemented with Ampicillin. GFP-nanotrap refers to the camelid-derived nanobody specific for GFP (Kubala *et al*, 2010). The expression and lysis of cells were carried out as described above. The supernatant containing GFP nano trap was first purified by affinity chromatography using Talon resin equilibrated with 50 mM Tris (pH 8.0), 500 mM NaCl, 10 % Glycerol and protein bound to the column was eluted using elution buffer containing 50 mM Tris (pH 8.0), 500 mM NaCl, 10 % Glycerol, 300 mM Imidazole. The fractions containing GFP-nanotrap was combined, concentrated and loaded on to a Superdex 75 (16/60) column equilibrated with 50 mM Hepes (pH 7.5), 200 mM NaCl, 0.5 mM TCEP (tris(2- carboxyethyl)phosphine). The fractions containing the protein were analysed by SDS PAGE, combined, and concentrated to the desired concentration. Rat Sx1a (Sx11-261-His), Munc18-1-His and Munc18-1^ι1317-333^ were also expressed and purified to homogeneity as described Hu et al (Hu *et al*., 2011).

### Cell culture and Transfection

Neurosecretory cell line, Pheochromocytoma cells (PC12) and Munc18-1/ 2 double knockout PC12 cells (DKO-PC12)(Jiang *et al*., 2023), were cultured at 37 °C/5% CO2 in normal culture media (Dulbecco’s modified Eagle’s medium (High glucose, pyruvate, Gibco, Cat#11995), 7.2% Heat-inactivated Foetal Bovine Serum (Gibco), 7.2% Heat-inactivated Horse Serum (Gibco), 1x GlutaMAX^TM^ supplement (Gibco, Cat# 35050061)). Cells were transfected using Lipofectamine^TM^ LTX (Invitrogen, Cat#15338100) as per manufacturer’s instructions. For co-IP experiments PC12 cells 4 x 10^6^ cells were cultured in 10 cm culture dishes (TPP Techno Plastic Products AG) coated with 0.1 mg/mL Poly-D-Lysine (Sigma, P2636). For each condition 15.3 μg plasmid DNA was used per 10 cm dish and 2x 10 cm dishes were pooled for experiments. PC12 cells were transfected with either pEGFP-N1 (GFP Control), pcDNA3.1-N-eGFP hMint1 WT or pcDNA3.1-N-eGFP hMint1 DI/AA mutant, and 48hr post transfected cell pellets were collected for subsequent co-IP/GFP Trap. Munc18 DKO-PC12 cells were co-transfected with 1 μg pCI VAMP2-pHluorin and 1 μg of either pmEos3.2-N1 Munc18-1WT or pmEos3.2-N1 Munc18-1 R388A with 6.75 μL Lipofectamine^TM^ LTX with PLUS Reagent^TM^ (ThermoFisher, Cat# 15338-100) into 3.5 cm petri dishes as per the manufacturer’s instructions. Cells were re-plated after 24 h onto 0.1 mg/mL Poly-D-Lysine (SigmaAldrich, Cat#P2636-100MG) coated glass-bottom petri-dishes (Cellvis, Cat# D29-20-1.5-N) and imaged 48 hrs after transfection.

### Co-immunoprecipitation (Co-IP)

PC12 cells containing EGFP, GFP- hMint1 (WT), GFP hMint1 DI/AA were lysed on ice using buffer composed of 20 mM Hepes, 50 mM NaCl, 1% Triton, 1 mM DTT, DNAse and a tablet of protease inhibitor cocktail. To further enhance the Lysis, the lysate was aspirated through a small needle approximately 10 times. The lysate was then centrifuged at 17,000 *g* for 15 minutes to separate the cellular debris from the supernatant containing the soluble proteins. 50 *μl* of GFP nano trap coupled to NHS activated Sepharose 4 beads were added to each of the three supernatants containing GFP, GFP-hMint1 WT and GFP-hMint1 DI/AA. The supernatant-bead mixtures were incubated for 2 hours at 4 °C while shaking, and then the beads were spun down by at 5000 *g* for 2 minutes to remove the unbound proteins. The bead samples were washed three times using the lysis buffer, and 50 *μl* of SDS sample buffer was added to each sample. The beads, containing the immunoprecipitated proteins were boiled at 95 °C for 5 minutes to elute the bound proteins and resolved using Western immunoblotting. GFP and Munc18a proteins were detected using anti-GFP mouse and anti-Munc18a mouse as primary antibodies respectively and Goat anti-mouse antibody as the secondary antibody. The final imaging was performed using ECL and Odyssey infrared imaging system (LI-COR).

### Total Internal Reflection Fluorescence Microscopy (TIRFM) and cell footprint analysis

For live cell TIRF imaging, transfected Munc18 DKO-PC12 cells were imaged in glass-bottom dishes bathed in Buffer A (145 mM NaCl, 5 mM KCl, 1.2 mM Na2HPO4, 10 mM D-glucose, 20 mM HEPES, pH 7.4) and imaged immediately before and following 2 mM BaCl2 stimulation, to elicit vesicle fusion (Papadopulos *et al*, 2013). Dishes were imaged on an iLas2 Microscope (Roper Scientific) equipped with a Nikon CFI Apo TIRF 100×/1.49 NA oil-immersion objective, and an Evolve 512 Delta EMCCD camera (Photometrics) and Metamorph Imaging Software version 7.7.8. Cells were imaged at 10 Hz (100 ms acquisition), for 3000 frames (300 seconds) at 37°C/5% CO2, and 30% of the initial 491 laser power in TIRF.

### Vesicle Fusion Assays in Munc18 DKO cells

To assess vesicle fluorescence over time a custom Python 3.8 pipeline was developed. For a typical 3000 frame TIRF acquisition, the data was read into a Python z-stack and divided into 100 frame intervals. The fluorescence at each interval was averaged, and fluorescent puncta identified at pixel resolution (where 1 pixel = 106 nm) using the Laplacian of Gaussian functionality of Python OpenCV. The puncta were used to populate a 3D [x,y,t] array. 3D DBSCAN (scikit-learn) was used to identify clusters of puncta which were within 1 pixel spatially and 2 pixels temporally. The time corresponding to the disappearance of each cluster was used as the indicator of a completed fusion event. Ongoing clusters that had not disappeared by the end of the acquisition were not considered for further analysis.

### Statistical Analysis of vesicle fusion events in Munc18 DKO cells

Unless otherwise stated, values are represented as mean±SEM. The tests used for statistical analysis are indicated in the respective figure legends. Non-parametric Mann Whitney U test was used to compare two groups of for non-normally distributed data. Comparisons of the same cells analysed before and after stimulation was analysed by a paired statistical test. Data were considered significant at P < 0.05.

### Isothermal titration calorimetry (ITC)

For ITC, all peptides were weighed and initially dissolved in the working buffer to make a stock concentration of 2 mM. ITC experiments measuring Munc18-1 binding to Mint1 peptides and Sx1a were carried out on Microcal iTC200 at 25^ο^C. GST-Mint1 and all the other proteins used in the experiments were buffer exchanged into 50 mM Tris, (pH 8.0), 200 mM NaCl by gel filtration prior to ITC. GST-Mint1 (1 mM), Sx1a (1 mM) or Mint1 synthetic peptide (0.7 mM) were titrated into Munc18- 1-His (50 μM). Mint1 PTB domain binding to the APP and TJAP1 peptides was performed in 50 mM HEPES, 200 mM NaCl, 0.5 mM TCEP and 5% glycerol at 25^ο^C with 1 mM peptides titrated into 50 μM Mint1 PTB domain. The binding parameters, equilibrium constant *K*a (1/Kd), stoichiometry (n) and the enthalpy *(ι1H*) were determined by processing the ITC data using ORIGIN 7.0 software. The equation, *ι1G* = *ι1H* – T*ι1S* was used to calculate the Gibbs free energy (*ι1G*) and all the experiments were performed in triplicate or duplicate to calculate the standard error of the mean (SEM) for thermodynamic parameters.

### Pull down assays

Pull down assays were carried out using GST-Mint1 and Munc18-1-His. 0.5 nmol of GST-Mint1 was mixed with 1 nmol of Munc18-1-His in 500 μL of pull down buffer (50 mM Tris pH 8.0, 200 mM NaCl, 1mM DTT, 0.1% IGEPAL) and incubated for 30 min on a rotating holder at 4 ^ο^C. The protein mixture was then centrifuged at full speed for 5 min and 50 μL of glutathione Sepharose resin pre-equilibrated in pull down buffer was added. The protein mix with the resin was incubated further 30 min at 4 ^ο^C on a rotating holder and at the end of the incubation, the beads were spun down at 5000 rpm for 30 sec. The supernatant containing the unbound protein is pipetted off and the beads with the bound protein was washed 4 times with 1 mL of pull down buffer. 50 μL of SDS sample buffer was added to the beads and analysed by SDS-PAGE for bound proteins.

### Protein structural prediction, modelling and visualisation

All protein models were generated using AlphaFold2 Multimer (Evans *et al*., 2022; Jumper *et al*., 2021) implemented in the Colabfold interface available on the Google Colab platform (Mirdita *et al*., 2022). For each modelling experiment ColabFold was executed using default settings where multiple sequence alignments were generated with MMseqs2 (Mirdita *et al*, 2019). For all final models displayed in this manuscript, structural relaxation of peptide geometry was performed with AMBER (Hornak *et al*, 2006). For all modelling experiments, we assessed (i) the prediction confidence measures (pLDDT and interfacial iPTM scores), (ii) the plots of the predicted alignment errors (PAE) and (iii) backbone alignments of the final structures. For modelling of the complex between Munc18- 1 and Mint1 we initially predicted the complex between full-length proteins and identified a high-confidence binding sequence in the N-terminal region of Mint1 that correlated precisely with the binding motif identified in our biochemical experiments. Based on this initial model and our biochemical mapping of the minimal Mint1 sequence for Munc18-1 binding we performed multiple independent predictions using a shorter peptide region, combined with AMBER energy minimization to optimize amino acid stereochemistry.

For interactome-wide analysis of other protein-protein interactors of Mint1 and Mint2 we obtained a list of putative interactors from the BioGRID repository (Oughtred *et al*., 2021), and predicted whether they formed direct complexes using the ColabFold Batch implementation of AlphaFold2 (Mirdita *et al*., 2022) (**Table S1; Table S2**). To assign a direct ‘interactor’ from these *in silico* analyses we used an approach similar to recent work by Sifri and colleagues (Sifri *et al*., 2023). We initially assessed both the AlphaFold2-derived interfacial PTM (iPTM) score and the resultant PAE graphs which provide confidence metrics for the interactions between the proteins. For those with promising scores we also examined the predicted structures in PyMOL to assess whether interacting regions involved the expected complementary hydrophobic, polar, and electrostatic contacts. To conserve GPU resource allocation, we initially generated three separate predictions of each putative complex in AlphaFold2 in unsupervised batch mode. We found that a minimum average iPTM score of ∼0.3 combined with a strong signal in the PAE plots for inter-molecular structural correlation typically provided a useful indicator of a complex that was suitable for further assessment. In these cases, we subsequently ran at least three modelling experiments focusing on the specific domains of Mint1/Mint2 and the putative interactors that were predicted to interact with each other, to assess whether multiple predictions resulted in physically plausible structures that consistently aligned with each other in PyMOL (**Table S1; Table S2**). Sequence conservation across PDB models was mapped with Consurf (Ashkenazy *et al*, 2016). All structural images were made with Pymol (Schrodinger, USA; https://pymol.org/2/).

## Acknowledgements

This work was supported by grants from the National Health and Medical Research Council of Australia (NHMRC) (APP1138083 and APP1139316) to B.M.C and FAM. B.M.C is supported by an NHMRC Investigator Grant and previous Senior Research Fellowship (APP2016410 and APP1136021). F.A.M is supported by an NHMRC Senior Research Fellowship (APP1155794). T.P.W is supported by an NHMRC Ideas Grant (APP2010901) awarded to T.P.W and F.A.M. We would like to thank Lionel Louiset, Ailisa Blum, Pranesh Padmanabhan and Jenny Martin for their assistance with this work, including initial mammalian construct testing, funding contributions and supervision of students.

## Author contributions

B.M.C. designed research with some input of F.A.M; S.W. and E.L. performed ITC analysis, pulldown assays and S.W. and B.M.C. performed AlphaFold2 modelling; S.W. and R.S.G performed the coimmunoprecipitation assays. R.S.G performed cell culture, plasmid design and cloning; A.M performed site directed mutagenesis with the help of R.S.G., cell culture and TIRFM with the help of A.T.B.. T.P.W designed the exocytic fusion assay Python pipeline and performed the analysis. A.J assisted with data processing and analysis. F.A.M. and B.M.C acquired funding and provided student supervision; and S.W., R.S.G., F.A.M. and B.M.C wrote the paper with input by all authors.

## Supplementary Information

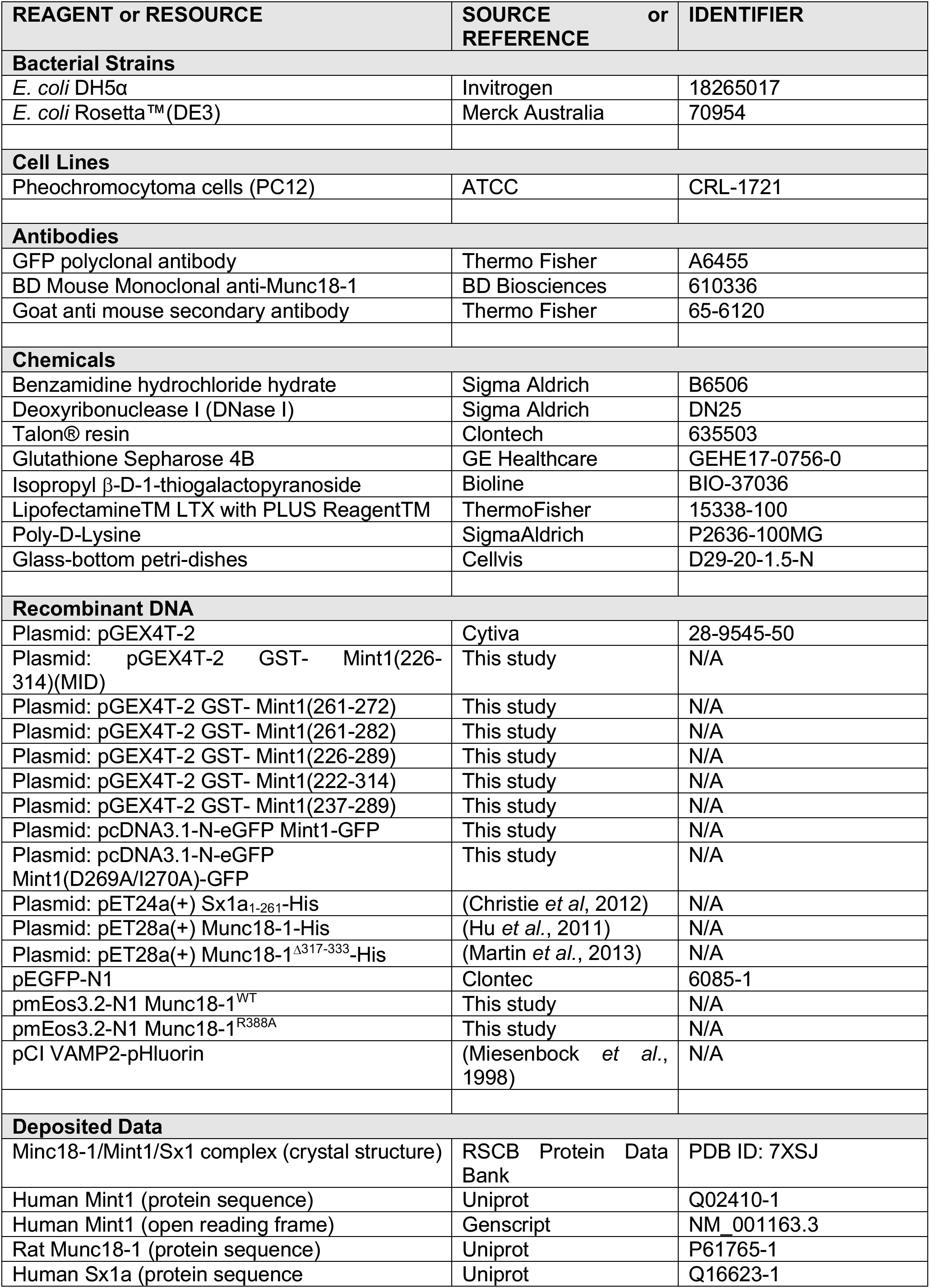
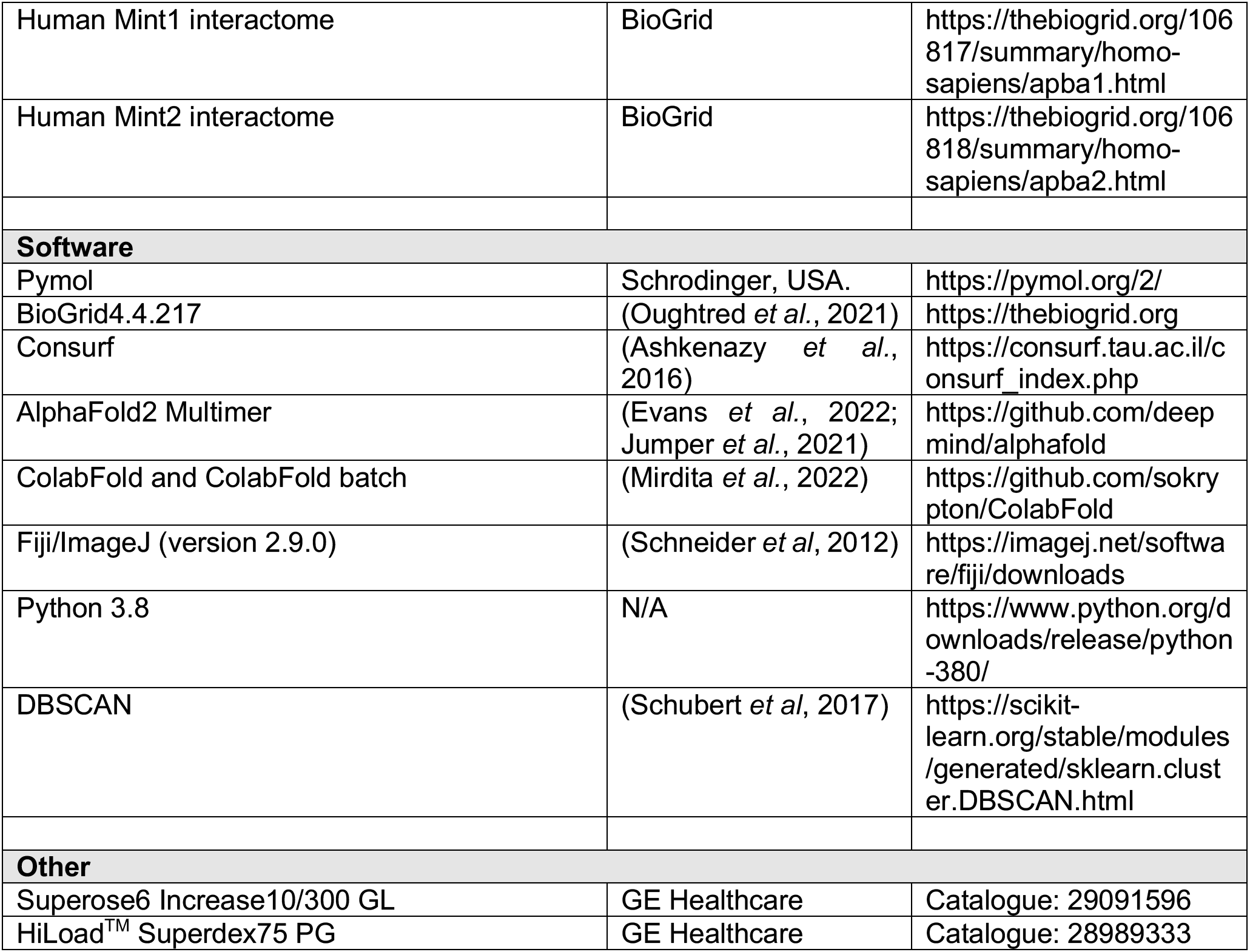
Resources Table.

**Table S1.**
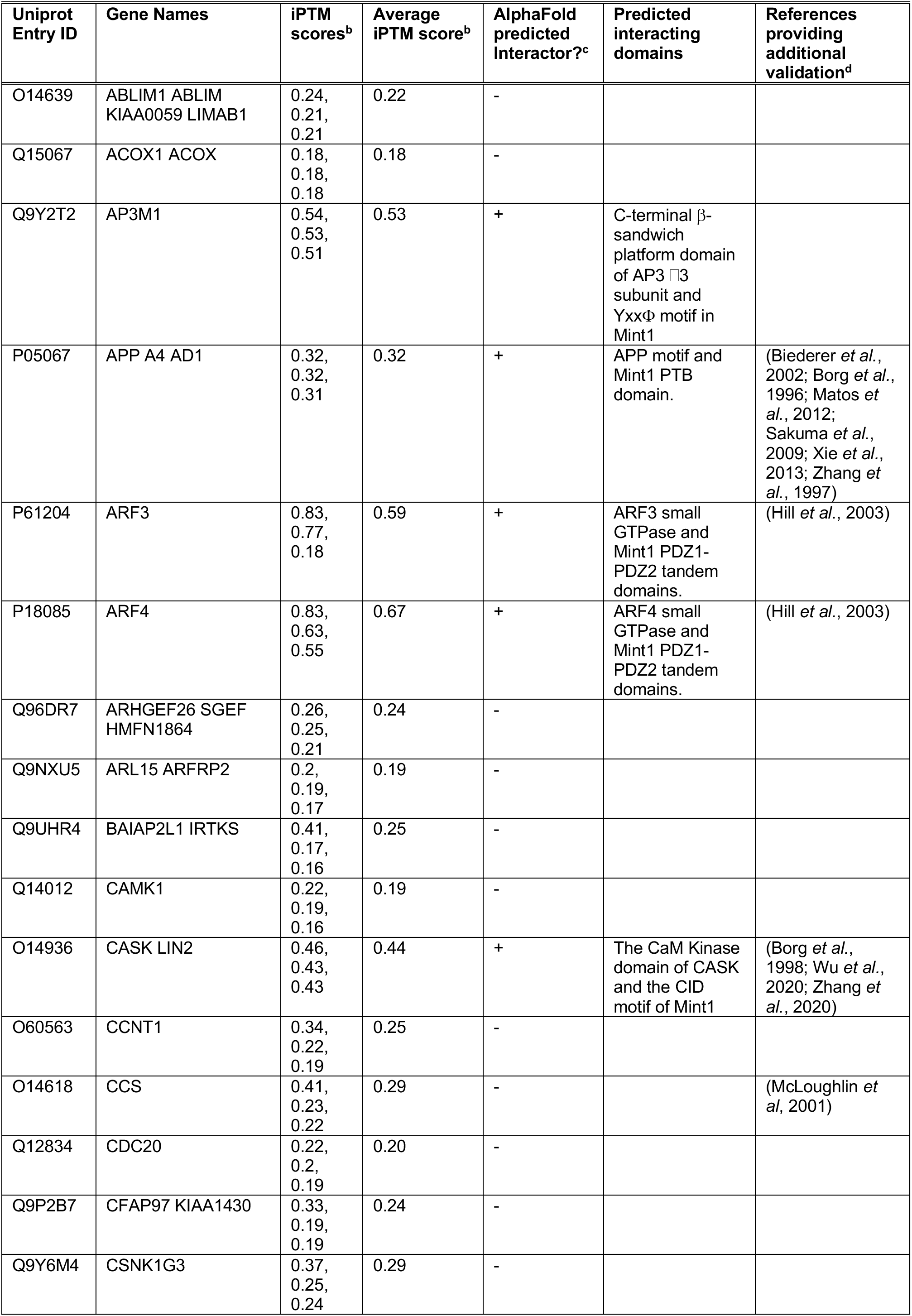
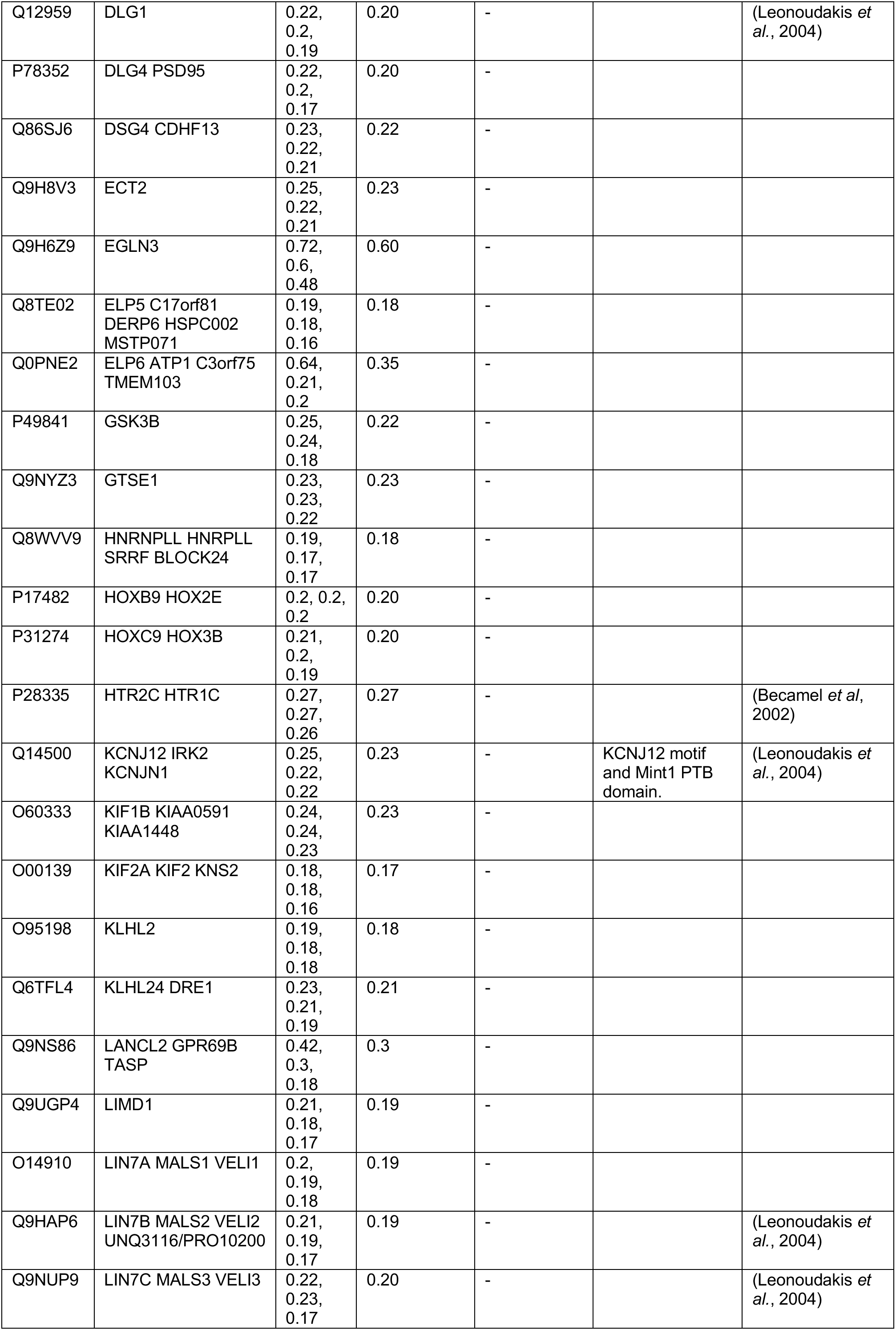
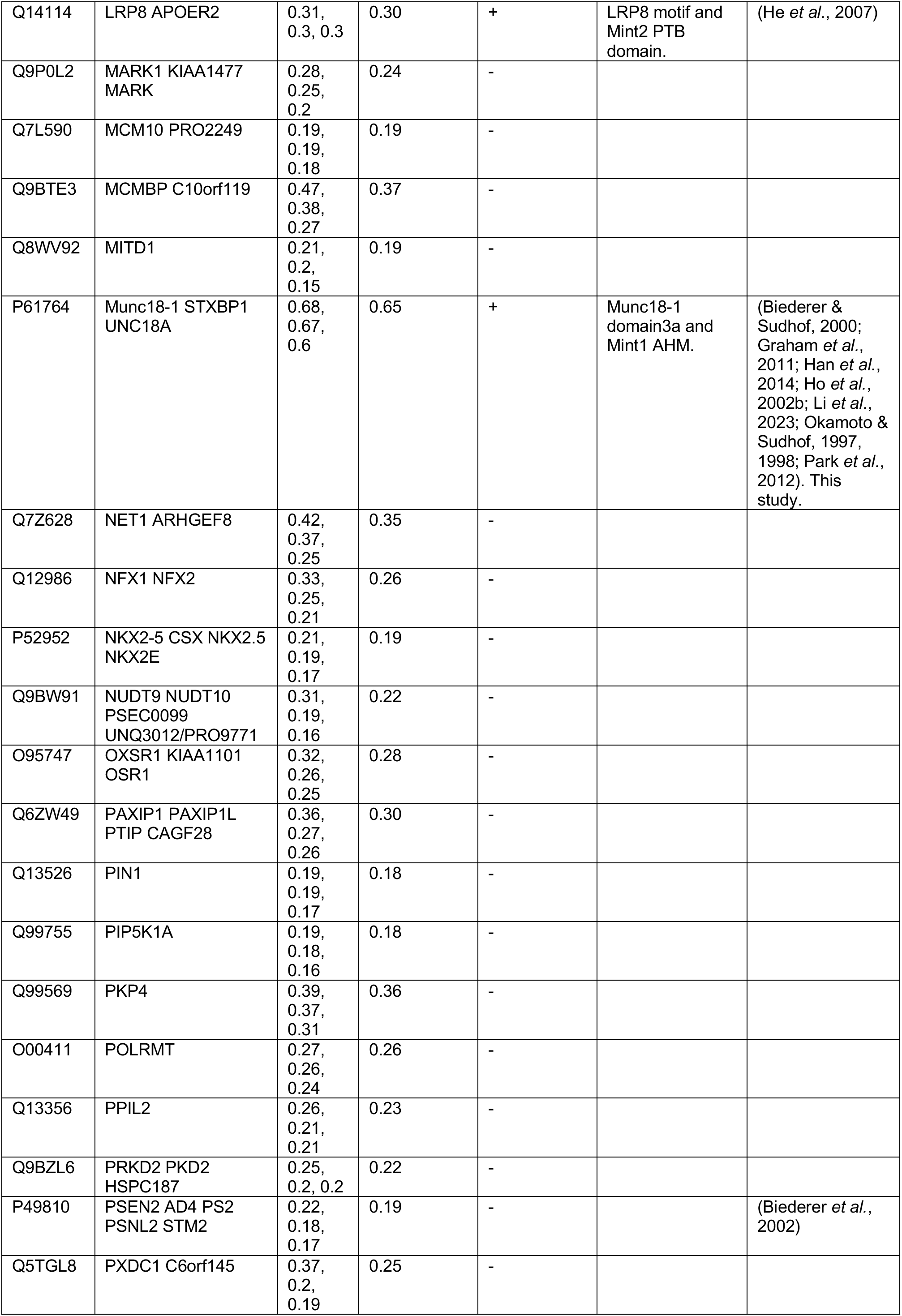
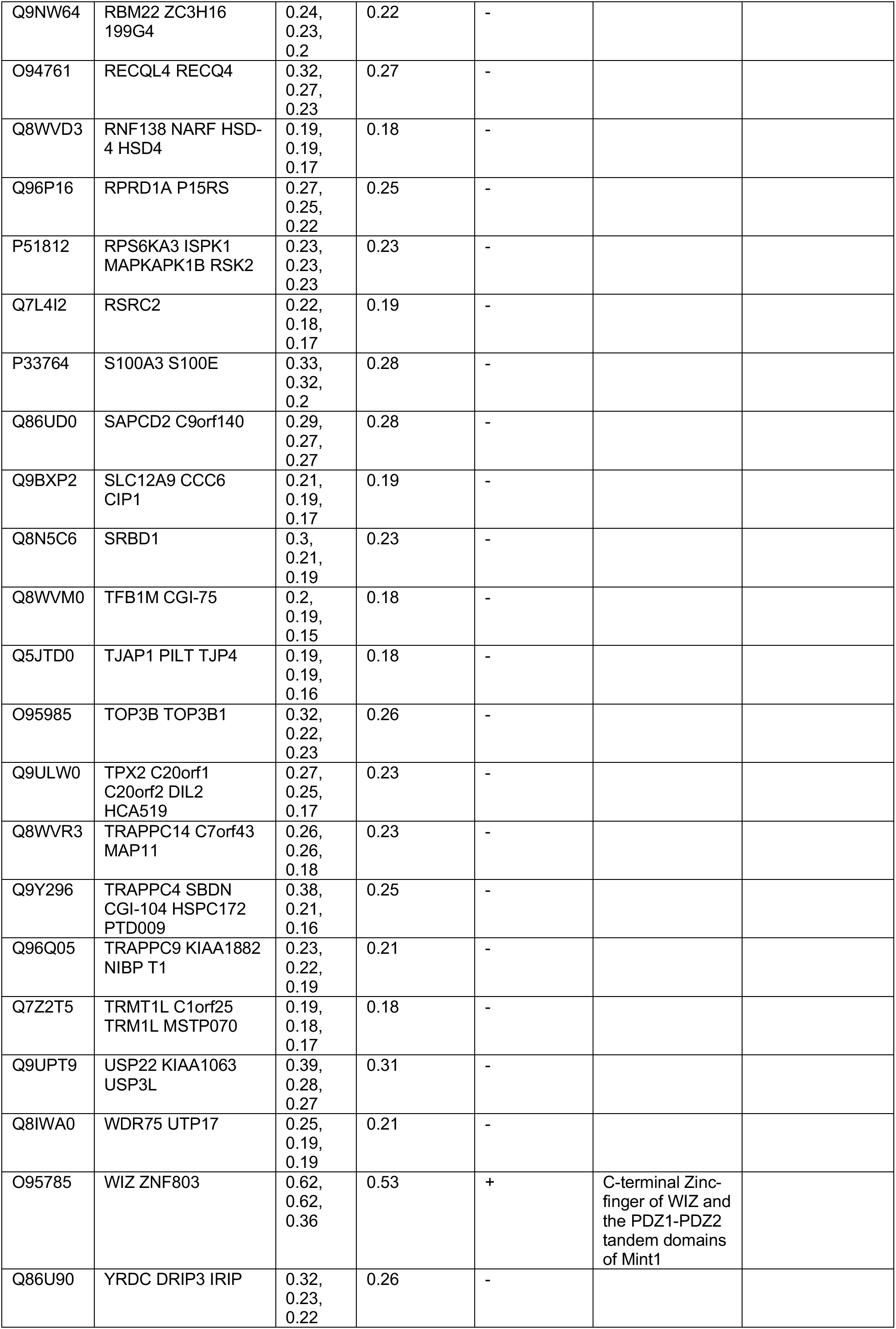
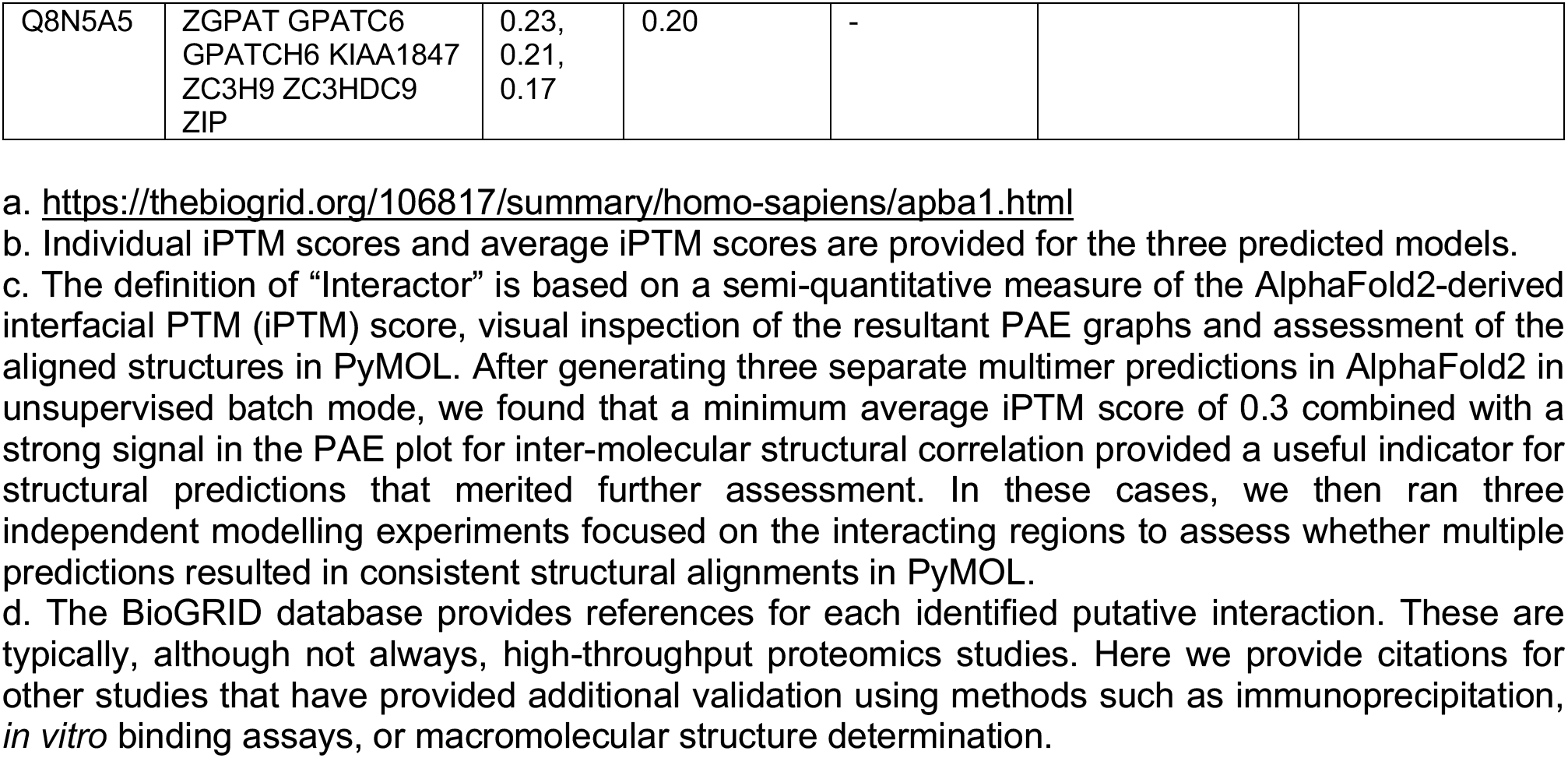
Putative interactors of human Mint1 from BioGRID^a^.

**Table S2.** Putative interactors of human Mint2 from BioGRID^a^.

| Uniprot Entry ID | Gene Names | iPTM scores<sup>b</sup> | Average iPTM score<sup>b</sup> | AlphaFold predicted Interactor?<sup>c</sup> | Predicted interacting domains | References providing additional validation<sup>d</sup> |
| --- | --- | --- | --- | --- | --- | --- |
| Q6PD74 | AAGAB | 0.17, 0.16, 0.16 | 0.16 | - |  |  |
| Q9UBZ4 | APEX2<br>APE2<br>APEXL2<br>XTH2 | 0.23, 0.22, 0.18 | 0.21 | - |  |  |
| P05067 | APP A4 AD1 | 0.84, 0.83, 0.80 | 0.82 | + | APP motif and Mint2 PTB domain. | (Borg *et al.*, 1996; Matos *et al.*, 2012; Xie *et al.*, 2013; Zhang *et al.*, 1997) |
| P61204 | ARF3 | 0.84, 0.81, 0.81 | 0.82 | + | ARF3 small GTPase and Mint2 PDZ1-PDZ2 tandem domains. | (Hill *et al.*, 2003) |
| P18085 | ARF4 | 0.87, 0.85, 0.84 | 0.85 | + | ARF4 small GTPase and Mint2 PDZ1-PDZ2 tandem domains. | (Hill *et al.*, 2003) |
| Q7Z3C6 | ATG9A<br>APG9L1 | 0.32, 0.3, 0.28 | 0.30 | - |  |  |
| O15155 | BET1 | 0.22, 0.21, 0.15 | 0.19 | - |  |  |
| Q9Y297 | BTRC<br>BTRCP<br>FBW1A<br>FBXW1A | 0.37, 0.32, 0.15 | 0.28 | - |  |  |
| A6NLJ0 | C2CD4B<br>FAM148B<br>NLF2 | 0.27, 0.2, 0.18 | 0.22 | - |  |  |
| O94985 | CLSTN1<br>CS1<br>KIAA0911 | 0.34, 0.34, 0.33 | 0.34 | + | CLSTN1 motif and Mint2 PTB domain. | (Araki *et al.*, 2003) |
| Q9GZR7 | DDX24 | 0.19, 0.19, 0.19 | 0.19 | - |  |  |
| Q09019 | DMWD DM9 | 0.41, 0.4, 0.29 | 0.37 | - |  |  |
| Q9UKB1 | FBXW11<br>BTRCP2<br>FBW1B<br>FBXW1B<br>KIAA0696 | 0.33, 0.19, 0.16 | 0.23 | - |  |  |
| A4D1S0 | KLRG2<br>CLEC15B | 0.19, 0.18, 0.16 | 0.18 | - |  |  |
| Q07954 | LRP1 A2MR<br>APR | 0.82, 0.82, 0.78 | 0.81 | + | LRP1 motif and Mint2 PTB domain. | (Gotthardt *et al.*, 2000) |
| P98164 | LRP2 | 0.61, 0.59, 0.55 | 0.59 | + | LRP2 motif and Mint2 PTB domain. | (Gotthardt *et al.*, 2000) |
| Q14114 | LRP8<br>APOER2 | 0.81, 0.77,<br>0.34 | 0.64 | + | LRP8 motif<br>and Mint2<br>PTB<br>domain. | (Gotthardt *et al.*, 2000) |
| Q8N7X4 | MAGEB6 | 0.38, 0.34,<br>0.33 | 0.35 | - |  |  |
| Q16659 | MAPK6<br>ERK3<br>PRKM6 | 0.26, 0.26,<br>0.19 | 0.24 | - |  |  |
| P61764 | Munc18-1<br>STXBP1<br>UNC18A | 0.67, 0.66,<br>0.63 | 0.65 | + | Munc18-1<br>domain3a<br>and Mint2<br>AHM. | (Biederer &<br>Sudhof, 2000;<br>Graham *et al.*,<br>2011; Han *et al.*,<br>2014; Ho<br>*et al.*, 2002b;<br>Li *et al.*, 2023;<br>Okamoto &<br>Sudhof, 1997,<br>1998; Park *et al.*,<br>2012).<br>This study. |
| Q96P71 | NECAB3<br>APBA2BP<br>NIP1 SYTIP2<br>XB51 | 0.16, 0.16,<br>0.14 | 0.15 | - |  | (Lee *et al.*,<br>2000) |
| Q13393 | PLD1 | 0.27, 0.22,<br>0.2 | 0.23 | - |  |  |
| P49810 | PSEN2 AD4<br>PS2 PSNL2<br>STM2 | 0.16, 0.15,<br>0.15 | 0.15 | - |  | (Biederer *et al.*,<br>2002) |
| O94761 | RECQL4<br>RECQ4 | 0.28, 0.28,<br>0.26 | 0.27 | - |  |  |
| Q04206 | RELA<br>NFKB3 | 0.23, 0.18,<br>0.17 | 0.19 | - |  | (Tomita *et al.*,<br>2000) |
| Q9NVW2 | RLIM RNF12 | 0.22, 0.2,<br>0.19 | 0.20 | - |  |  |
| Q68DV7 | RNF43 | 0.32, 0.28,<br>0.25 | 0.28 | - |  |  |
| O15431 | SLC31A1<br>COPT1<br>CTR1 | 0.27, 0.2,<br>0.18 | 0.22 | - |  |  |
| Q9HB58 | SP110 | 0.17, 0.17,<br>0.17 | 0.17 | - |  |  |
| Q8N9I0 | SYT2 | 0.17, 0.15,<br>0.15 | 0.16 | - |  |  |
| Q5JTD0 | TJAP1 PILT<br>TJP4 | 0.40, 0.31,<br>0.29 | 0.33 | + | TJAP1 N-terminal<br>sequence<br>and Mint2<br>PTB<br>domain in<br>non-canonical<br>site |  |
| Q8NFZ5 | TNIP2 ABIN2<br>FLIP1 | 0.16, 0.15,<br>0.14 | 0.15 | - |  |  |
| Q8WV44 | TRIM41<br>RINCK | 0.19, 0.17,<br>0.17 | 0.18 | - |  |  |
| Q9Y2K6 | USP20<br>KIAA1003<br>LSFR3A<br>VDU2 | 0.32, 0.28,<br>0.24 | 0.28 | - |  |  |
| Q8TEY7 | USP33<br>KIAA1097<br>VDU1 | 0.29, 0.27,<br>0.25 | 0.27 | - |  |  |
| Q6AHZ1 | ZNF518A<br>KIAA0335<br>ZNF518 | 0.23, 0.23,<br>0.16 | 0.21 | - |  |  |
| Q6DN90 | IQSEC1<br>ARFGEP100<br>BRAG2<br>KIAA0763 | 0.79, 0.78,<br>0.77 | 0.77 | + | IQSEC1<br>PH domain<br>and<br>peptide<br>sequence<br>in Mint2 N-terminal<br>region |  |
| Q5JU85 | IQSEC2<br>KIAA0522 | 0.66, 0.65,<br>0.59 | 0.65 | + | IQSEC2<br>PH domain<br>and<br>peptide<br>sequence<br>in Mint2 N-terminal<br>region |  |
| Q5VYV7 | SLX4IP<br>C20orf94 | 0.19, 0.19,<br>0.16 | 0.18 | - |  |  |

a. https://thebiogrid.org/106817/summary/homo-sapiens/apba1.html
b. Individual iPTM scores and average iPTM scores are provided for the three predicted models.
c. The definition of “Interactor” is based on a semi-quantitative measure of the AlphaFold2-derived interfacial PTM (iPTM) score, visual inspection of the resultant PAE graphs and assessment of the aligned structures in PyMOL. After generating three separate multimer predictions in AlphaFold2 in unsupervised batch mode, we found that a minimum average iPTM score of 0.3 combined with a strong signal in the PAE plot for inter-molecular structural correlation provided a useful indicator for structural predictions that merited further assessment. In these cases, we then ran three independent modelling experiments focused on the interacting regions to assess whether multiple predictions resulted in consistent structural alignments in PyMOL.
d. The BioGRID database provides references for each identified putative interaction. These are typically, although not always, high-throughput proteomics studies. Here we provide citations for other studies that have provided additional validation using methods such as immunoprecipitation, *in vitro* binding assays, or macromolecular structure determination.

**Figure S1.**
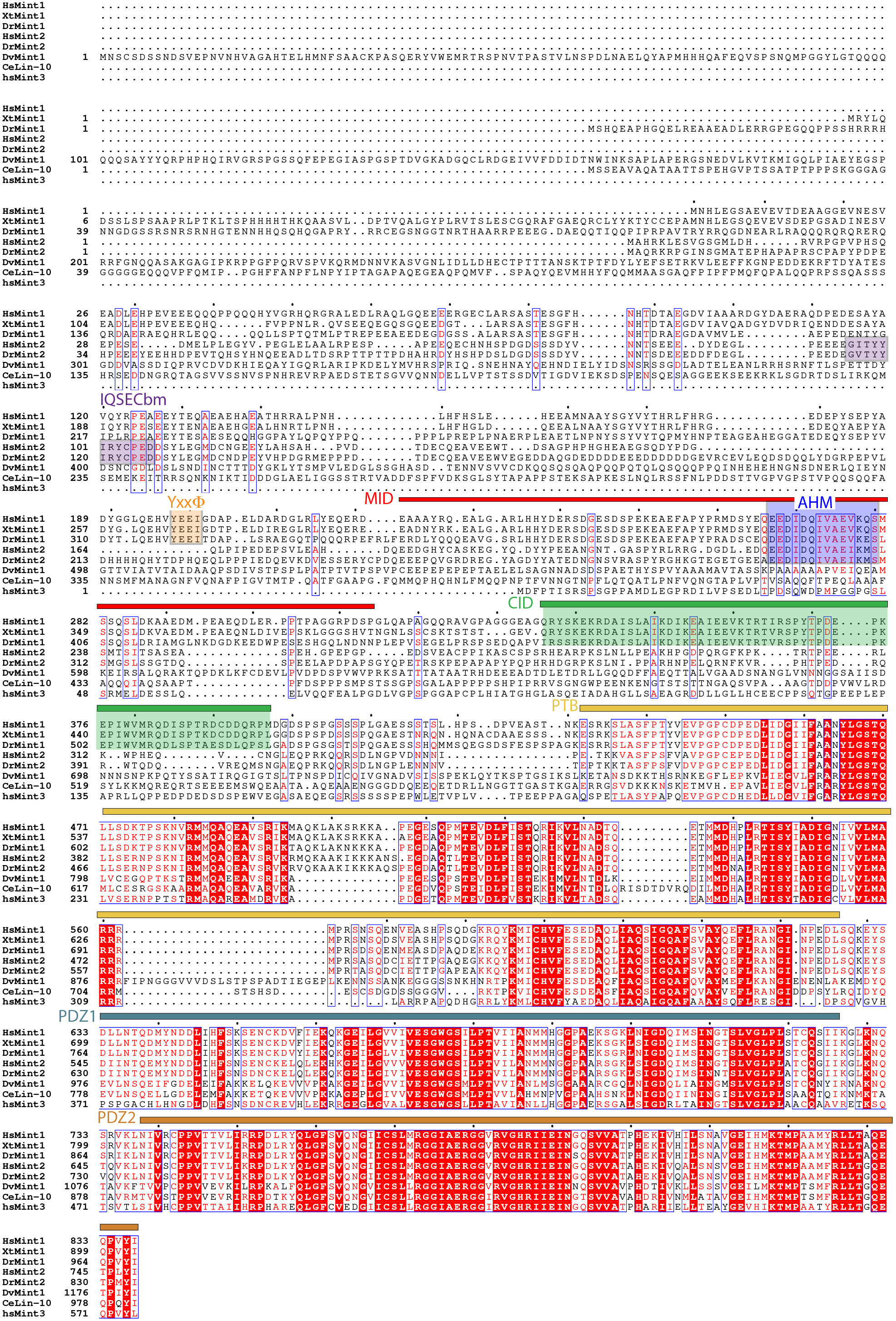
Sequence alignment of Mint proteins. (Related to Fig. 1) **(A)** Sequence alignment Mint1, Mint2 and Mint3. Alignment was performed with Multalin (Corpet *et al*, 1999) and rendered with ESPript3.0 (Robert & Gouet, 2014). Hs, *Homo sapiens*. Xt, *Xenopus tropicalis*. Dr, *Danio rerio*. Ce, *Caenorhabditis elegans*. Dv, *Drosophila virilis*.

**Figure S2.**
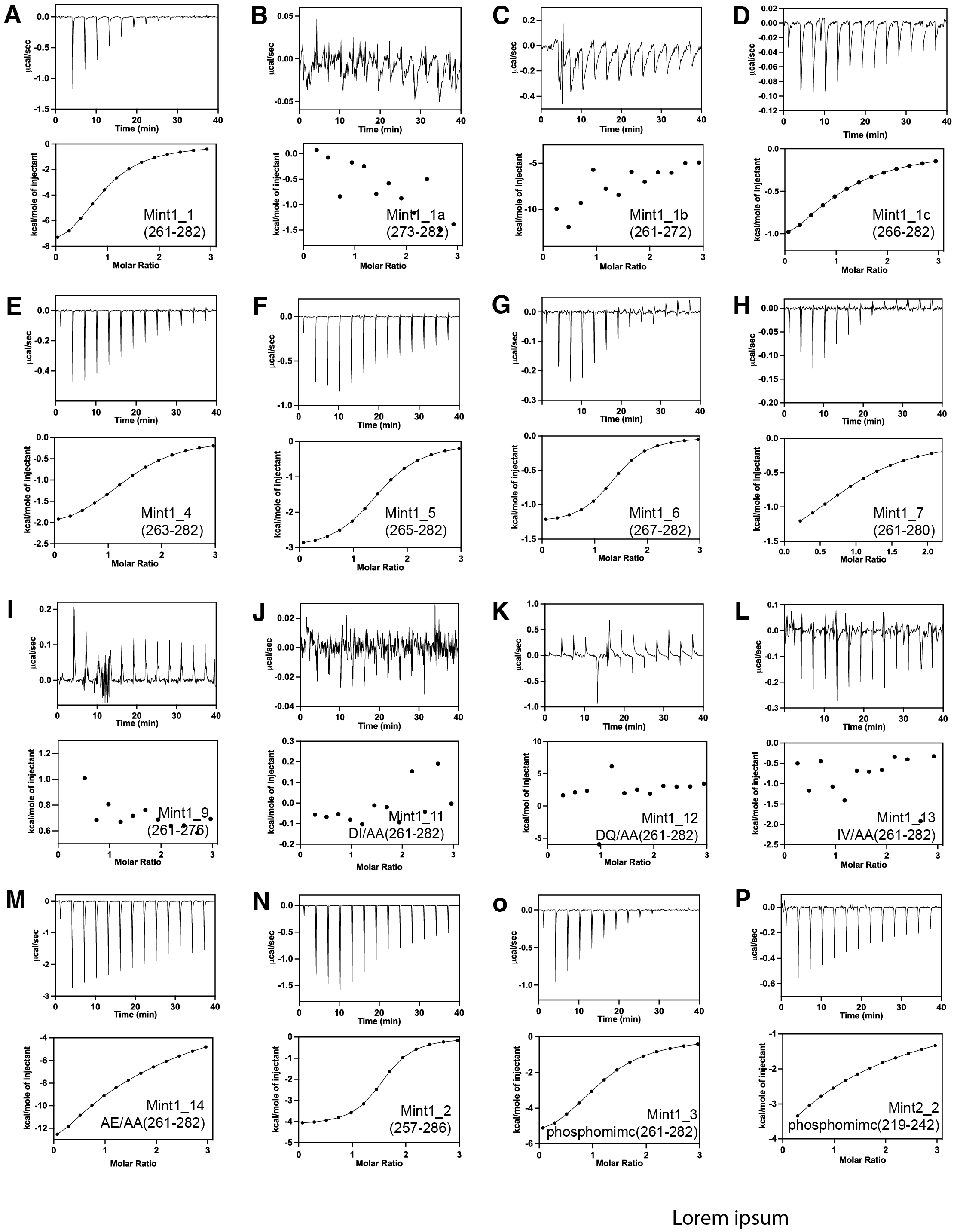
ITC experiments of Munc18-1 binding to various Mint peptides. (Related to Fig. 2) Example ITC experiments are shown for each of the Mint peptides binding to Munc18-1 as described in Fig. 2B and Table 1. (**A**) Mint 1_1 (261-282) (**B**) Mint 1_1a (273-282) (**C**) Mint 1_1b (261-272) (**D**) Mint 1_1c (266-282) (**E**) Mint 1_4 (263-282) (**F**) Mint 1_5 (265-282) (**G**) Mint 1_6 (267-282) (**H**) Mint 1_7 (261-280) (**I**) Mint 1_9 (261-276) (**J**) Mint 1_11 DI /AA (261-282 (**K**) Mint 1-12 DQ /AA (261-282) (**L**) Mint 1_13 IV/AA(261-282) (**M**) Mint 1_14 AE/AA(261-282) (**N**) Mint 1_2 (257-286) (**O**) Mint 1_3 phosphomimetic (261-282) (**P**) Mint 2_2 phosphomimetic (219-242). **Table 1** shows the binding affinities (Kd) of the above Mint 1 peptides obtained by integrating and normalising data fit with 1:1 ratio binding model. The Kd for Mint 1-1, 1a,1b and 1c are given as a mean of at least 2 independent experiments (n=2) and for all the other peptides Kds have been calculated from a single experiment (n=1). The peptides for which binding was not detected are shown as ‘nb,’ abbreviating ‘No Binding was detected”.

**Figure S3.**
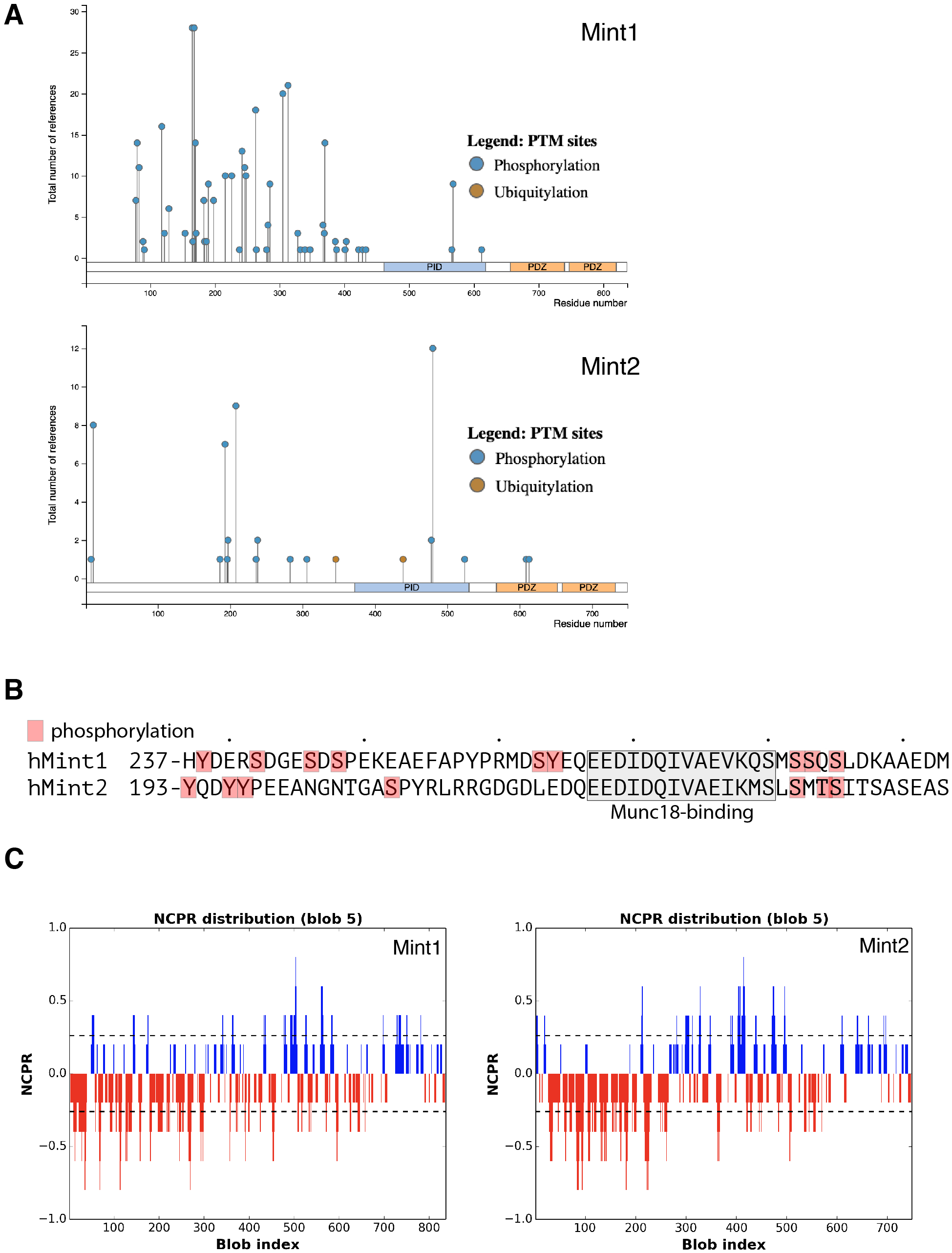
Post-translational modifications and sequence charge distribution of human Mint1 and Mint2. (Related to Fig. 2) (**A**) Sites of phosphorylation and ubiquitination of human Mint1 and Mint2 experimentally observed and reported in PhosphoSitePlus (Hornbeck *et al*., 2015). **(B)** Phosphorylation sites in the human Mint1 and Mint2 N-terminal regions documented in PhosphoSitePlus (Hornbeck *et al*., 2015). Phosphomimetic mutations in regions adjacent to the Munc18-1 binding sequence do not influence affinity (Fig. 2B; **Table 1**). (**C**) Plot of the charge distribution versus the sequence of Mint1 and Mint2 shows that the N-terminal disordered sequences have a high net-positive charge. Plots were made with CIDER (Holehouse *et al*, 2015).

**Figure S4.**
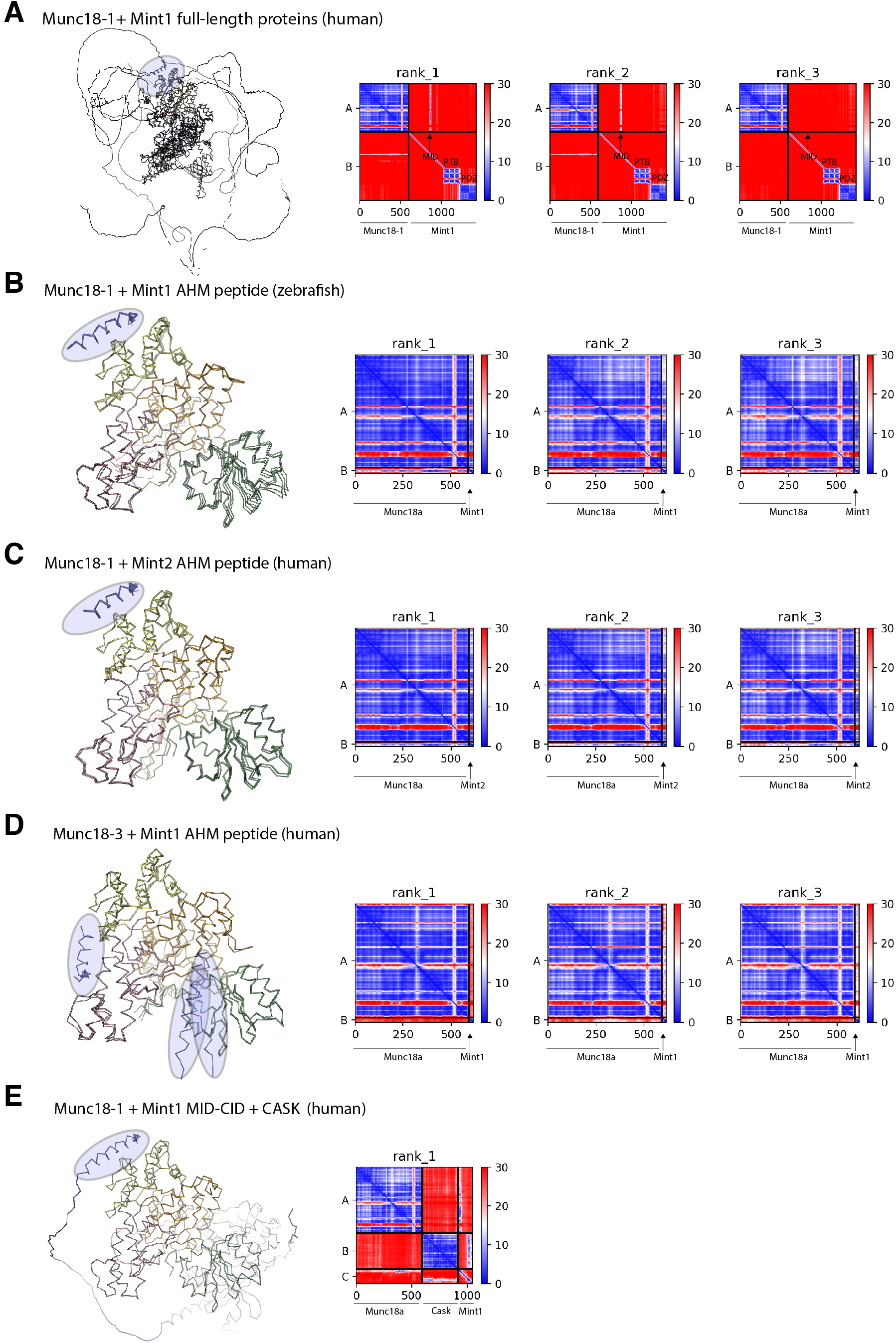
AlphaFold2 modelling of Munc18 interactions with Mint1. (Related to Fig. 3) **(A)** Overlaid top three ranked models of human Mint1 bound to Munc18-1 from ColabFold shown in backbone ribbon representation. Mint1 is consistently modelled to bind Munc18-1 domain 3b via a short acidic α-helical motif (AHM) within its unstructured N-terminal domain (highlighted in blue). On the right hand side the predicted alignment error (PAE) is plotted for each model. Signals in the off-diagonal regions indicate strong structural correlations between residues in the peptide with the Munc18-1 protein. (**B**) Overlaid top three ranked models of the zebrafish Mint1 AHM sequence bound to Munc18-1 predicted with ColabFold. Similar to the human structures, the AHM is consistently modelled in an α-helical structure associated with the Munc18-1 domain3b (highlighted in blue). (**C**) Overlaid top three ranked models of the human Mint2 AHM sequence bound to Munc18- 1 predicted with ColabFold. (**D**) Overlaid top three ranked models of the human Mint1 AHM sequence modelled with the non-binding Munc18-3 homologue. The AHM is modelled at several random positions (highlighted in blue). The PAE plots do not show evidence of significant interactions between the two proteins. (**E**) The top scoring model of the human Mint1 AHM and CID sequences bound to Munc18-1 and CASK using ColabFold. The AHM region is predicted in an identical binding site on Munc18-1 to the isolated peptide sequence (highlighted in blue). The CID sequence is predicted to bind to CASK in almost essentially the same conformation as the two previous crystal structures (Wu *et al*., 2020; Zhang *et al*., 2020).

**Figure S5.**
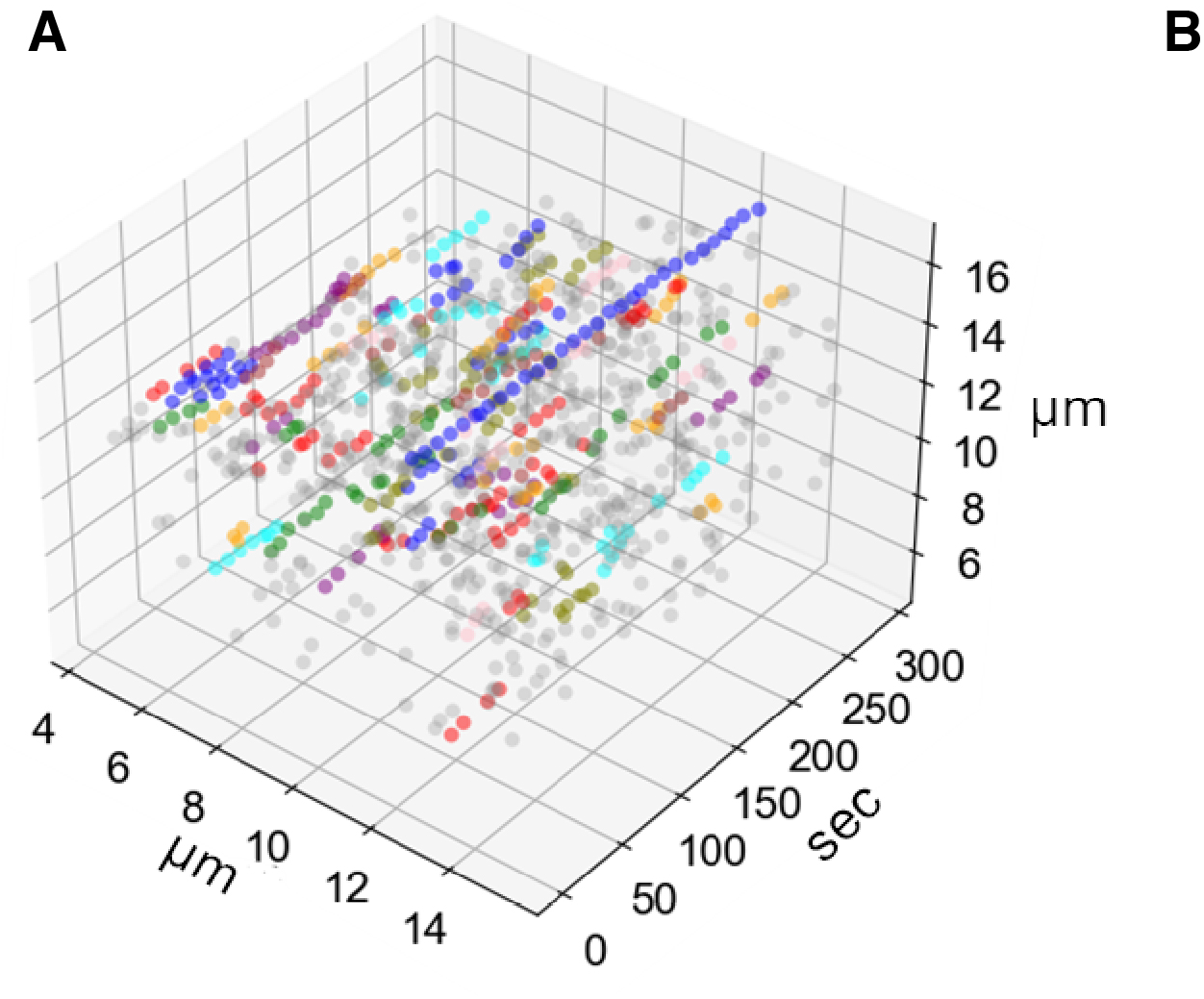
Analysis of cell footprint and fusion events in Munc18 DKO PC12 cells. Representative fluorescent release event visualisation. The imaging dataset was divided into 10 sec (100 frame) intervals and the fluorescence averaged. Vesicle fusions at each interval were determined by Laplacian of Gaussian and used to create a 3D [x,y,t] array. DBSCAN was used to determine clusters of fluorescent areas/vesicles which persisted in the same area over time. A release event was defined as the end of each cluster.

**Figure S6.**
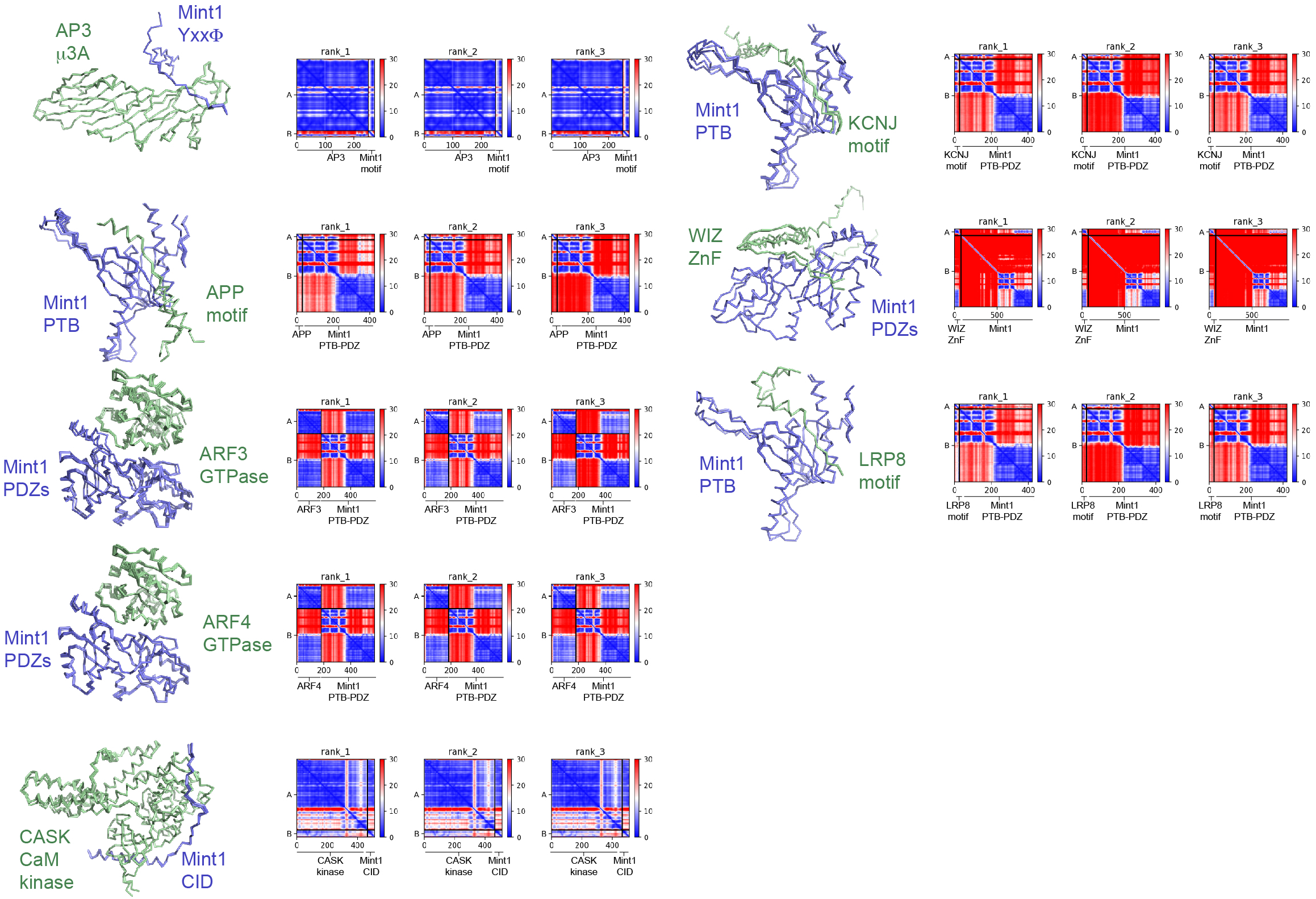
AlphaFold2 modelling of Mint1 interactions with proteins identified in BioGRID. (Related to Fig. 5 and 6) Proteins identified in BioGRID as putative interactors of Mint1 and showing reasonable binding in AlphaFold2 predictions. The left panels show the top three ranked structures overlaid in backbone ribbon representation. On the right hand side, the predicted alignment error (PAE) is plotted for each model. Signals in the off-diagonal regions indicate strong structural correlations between residues in the peptide with the Mint1 protein. These focused predictions were performed on specific Mint1 domains with the ligand regions initially identified in high-throughput predictions of the two full-length proteins.

**Figure S7.**
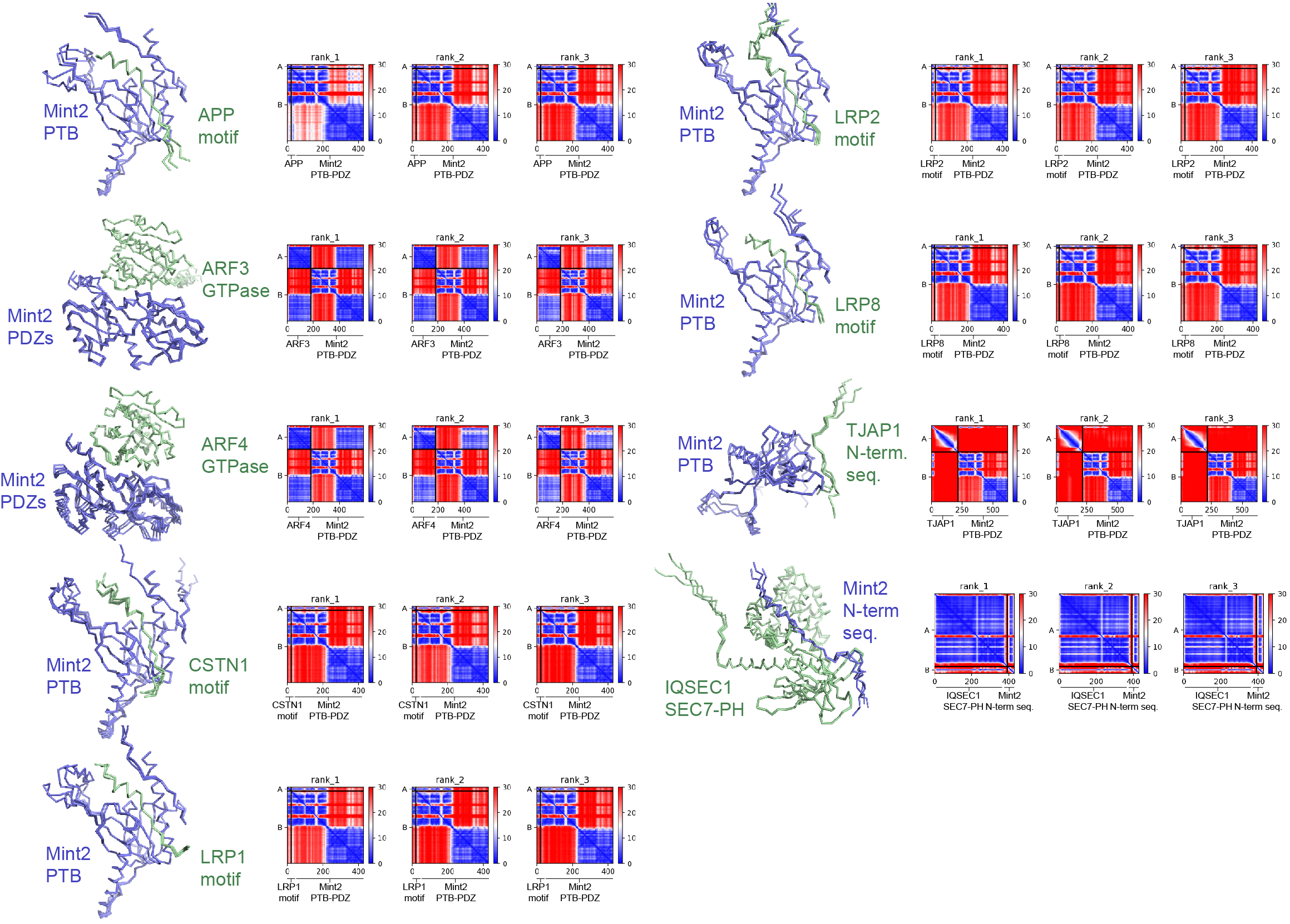
AlphaFold2 modelling of Mint2 interactions with proteins identified in BioGRID. (Related to Fig. 5 and 6) Proteins identified in BioGRID as putative interactors of Mint2 and showing reasonable binding in AlphaFold2 predictions. The left panels show the top three ranked structures overlaid in backbone ribbon representation. On the right hand side the predicted alignment error (PAE) is plotted for each model. Signals in the off-diagonal regions indicate strong structural correlations between residues in the peptide with the Mint2 protein. These focused predictions were performed on specific Mint2 domains with the ligand regions initially identified in high-throughput predictions of the two full-length proteins.

